# Probabilistic cell type assignment of single-cell transcriptomic data reveals spatiotemporal microenvironment dynamics in human cancers

**DOI:** 10.1101/521914

**Authors:** Allen W Zhang, Ciara O’Flanagan, Elizabeth A Chavez, Jamie LP Lim, Andrew McPherson, Matt Wiens, Pascale Walters, Tim Chan, Brittany Hewitson, Daniel Lai, Anja Mottok, Clementine Sarkozy, Lauren Chong, Tomohiro Aoki, Xuehai Wang, Andrew P Weng, Jessica N McAlpine, Samuel Aparicio, Christian Steidl, Kieran R Campbell, Sohrab P Shah

## Abstract

Single-cell RNA sequencing (scRNA-seq) has transformed biomedical research, enabling decomposition of complex tissues into disaggregated, functionally distinct cell types. For many applications, investigators wish to identify cell types with known marker genes. Typically, such cell type assignments are performed through unsupervised clustering followed by manual annotation based on these marker genes, or via “mapping” procedures to existing data. However, the manual interpretation required in the former case scales poorly to large datasets, which are also often prone to batch effects, while existing data for purified cell types must be available for the latter. Furthermore, unsupervised clustering can be error-prone, leading to under- and overclustering of the cell types of interest. To overcome these issues we present *CellAssign*, a probabilistic model that leverages prior knowledge of cell type marker genes to annotate scRNA-seq data into pre-defined and *de novo* cell types. CellAssign automates the process of assigning cells in a highly scalable manner across large datasets while simultaneously controlling for batch and patient effects. We demonstrate the analytical advantages of CellAssign through extensive simulations and exemplify realworld utility to profile the spatial dynamics of high-grade serous ovarian cancer and the temporal dynamics of follicular lymphoma. Our analysis reveals subclonal malignant phenotypes and points towards an evolutionary interplay between immune and cancer cell populations with cancer cells escaping immune recognition.

## 1 Introduction

Gene expression observed at the single-cell resolution in human tissues enables studying the cell type composition and dynamics of mixed cell populations in a variety of biological contexts, including cancer progression. Cell types inferred from single-cell RNA-seq (scRNA-seq) data are typically annotated in a two-step process, whereby cells are first clustered using unsupervised algorithms and then clusters are labeled with cell types according to aggregated cluster-level expression profiles [1]. A myriad of methods for unsupervised clustering of scRNA-seq have been proposed, such as SC3 [2], Seurat [3], PCAReduce [4], and PhenoGraph [5], along with studies evaluating their performance across a range of settings [6, 7]. However, clustering of low-dimensional projections may limit biological interpretability due to i) low-dimensional projections not encoding variation present in high-dimensional inputs [8] and ii) overclustering of populations that are not sufficiently variable.

Furthermore, even in the context of robust clustering which recapitulates biological cell states or classes, few principled methods for annotating clusters of cells into known cell types exist. In contrast to unsupervised statistical frameworks, this latter step is a supervised, or classification problem. Typical workflows employ differential expression analysis between clusters to manually classify cells according to highly differentially expressed markers, aided by recent databases linking cell types to canonical gene-based markers [9]. In situations where investigators wish to identify and quantify specific cell types of interest with known marker genes across multiple samples or replicates, such workflows can be cumbersome, and differences in clustering strategies can affect downstream interpretation [6]. Alternatively, cell types may be assigned by gating on marker gene expression, but this strategy is difficult to implement in practice as (i) gating is difficult for more than a few genes and relies on knowledge of marker gene expression levels and (ii) cells that fall outside these gates will not be assigned to any type, rather than being probabilistically assigned to the most likely cell type.

Another approach to cell type annotation is to leverage ground-truth single-cell transcriptomic data from labeled and purified cell types to establish robust profiles against which new data can be compared and classified. For example, scmap-cluster [10] calculates the medioid expression profile for each cell type in the known transcriptomic data, and then assigns input cells based on maximal correlation to those profiles. However, this approach requires existing scRNA-seq data for purified cell populations of interest. Given the technical effects associated with differences in experimental design and processing, expression profiles for reference populations may not be directly comparable to those for other single-cell RNA-seq experiments [11].

We assert that statistical cell type classification approaches leveraging prior knowledge in the literature (or from experiments) will be an effective complement to unsupervised approaches for quantitative decomposition of heterogeneous tissues from scRNA-seq data. Therefore, to address the analytical challenges inherent in both clustering and mapping approaches, we developed CellAssign, a scalable statistical framework that annotates and quantifies both known and *de novo* cell types in scRNA-seq data. CellAssign automates the process of annotation by encoding a set of *a priori* marker genes for each cell type. The statistical model then classifies the most likely cell type for each cell in the input data, using a marker gene matrix (cell type-by-gene). The model allows for flexible expression of marker genes, assuming that marker genes are more highly expressed in the cell types they define relative to others. Implemented in Google’s Tensorflow framework, CellAssign is highly scalable, capable of annotating thousands of cells in seconds while controlling for inter-batch, patient and site variability. We evaluated CellAssign across a range of simulation contexts and on ground truth data for FACS-purified H7 human embryonic stem cells (HSCs) at various differentiation stages [12], showing that CellAssign outperforms both clustering and correlation based methods—more readily discriminating closely related cell types—and is robust to errors in marker gene specification. In addition, we applied CellAssign to two novel datasets generated to profile spatiotemporal tumor microenvironment (TME) dynamics in human cancers. Using the CellAssign approach, we demonstrated tumor ‘ecosystem’ spatial diversity in untreated high-grade serous ovarian cancer through variable composition in stromal and immunologic cell types comprising the TME and variation in key pathways across malignant cell populations including immune evasion, epithelial-mesenchymal transition and hypoxia. Temporal dynamics were also exemplified using the CellAssign approach. We generated scRNA-seq libraries from matched diagnostic and relapsed pairs of follicular lymphoma samples, with one case having undergone histologic transformation to an aggressive lymphoma. We show compositional and phenotypic changes, including T-cell activation and HLA downregulation in cancer cells upon transformation, pointing towards an evolutionary interplay with cancer cells escaping immune recognition following transformation. In aggregate we conclude the CellAssign approach provides a robust new statistical framework through which disease dynamics in tissues comprised of mixed cell populations can be quantified and interpreted to ultimately uncover new properties and understanding of disease progression.

## 2 Results

### 2.1 CellAssign: probabilistic and automated cell type assignment

The CellAssign statistical framework (**Figure 1**) models observed gene expression for a heterogeneous cell population as a composite of multiple factors including cell type, library size, and batch. The inputs consist of raw single cell RNA-seq read counts and a marker gene set for each cell type of interest. Marker genes are assumed to be overexpressed in cell types where they are markers—not necessarily at similar levels—compared to those where they are not. Other experimental and biological covariates such as batch and patient-of-origin are optionally encoded in a standard design matrix. Using this information, CellAssign employs a hierarchical Bayesian statistical framework to determine the probability that each cell belongs to each of the modeled cell types, and estimates model parameters including the relative expression of marker genes in each cell type and the systematic effects of other covariates on marker gene expression patterns using an expectation-maximization inference algorithm. To prevent misclassification when unknown cell types (unspecified in the marker matrix) are present, CellAssign designates cells that do not belong to any provided cell type as ‘unassigned’. Detailed model specification, implementation and runtime performance are described in **Methods**.

**Figure 1.**
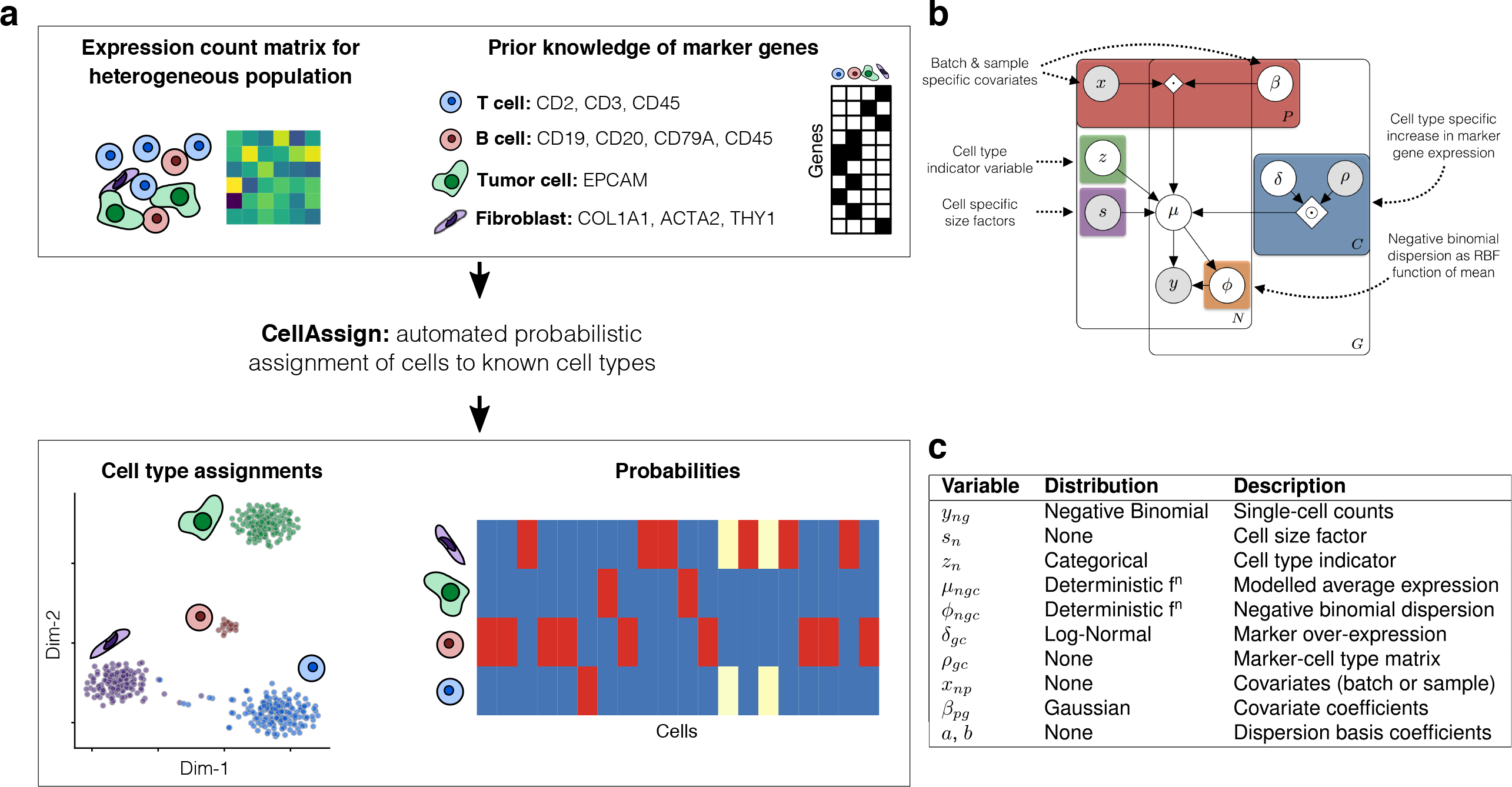
(**a**) Overview of CellAssign. CellAssign takes raw count data from a heterogeneous single-cell RNA-seq population, along with a set of known marker genes for various cell types under study. Using CellAssign for inference, each cell is probabilistically assigned to a given cell type without any need for manual annotation or intervention, accounting for any batch or sample-specific effects. (**b**) An overview of the CellAssign probabilistic graphical model. (**c**) The random variables and data that form the model, along with the distributional assumptions.

### 2.2 Performance of CellAssign relative to unsupervised clustering and supervised classification methods

We benchmarked CellAssign’s performance relative to standard workflows including unsupervised clustering followed by manual curation and methods that map cells to existing data from purified populations. Using an adapted version of the `splatter` model fitted to data for peripheral blood naïve CD8+ and CD4+ T cells, we simulated scRNA-seq data for multiple cell populations (**Methods**). Simulations were conducted across a wide range of values for differentially expressed gene fraction (0.05 to 0.45), to represent cellular mixtures of similar and distinct cell types. We then evaluated the performance of unsupervised (Seurat [3], SC3 [2], phenograph [13], densitycut [14], dynamicTreeCut [15]) and supervised (scmap-cluster [10], correlation-based [16]) methods (**Methods**). Half of the simulated cells (*n*=1000 training, *n*=1000 evaluation) were set aside exclusively for training the supervised methods. Marker genes for CellAssign were selected based on simulated log-fold change values and mean expression (**Methods**), and *maximum a posteriori* (MAP) cell type probability estimates were treated as deterministic cell type assignments. For all values of differentially expressed gene fraction, CellAssign performed better than alternative workflows in both accuracy and F1 score metrics (**Figure 2A**, **Supplemental Table 1**). Supervised methods generally performed better than unsupervised methods (**Figure 2A**). We then investigated the degree to which CellAssign’s performance was due to being provided with informative marker genes rather than transcriptome-wide data, repeating the analysis by providing other methods with exactly the same data as CellAssign. In this setting, CellAssign was still more accurate than the other methods (**Figure 2B**). Similar results were obtained on data simulated from parameter estimates fitted to B cells and CD8+ T cells (**Supplemental Figure 1A,B**, **Supplemental Table 1**). Moreover, CellAssign accurately inferred the relative expression of marker genes in each cell type (all *R* > 0.958; **Figure 2C**, **Supplemental Figure 1C**).

**Figure 2.**
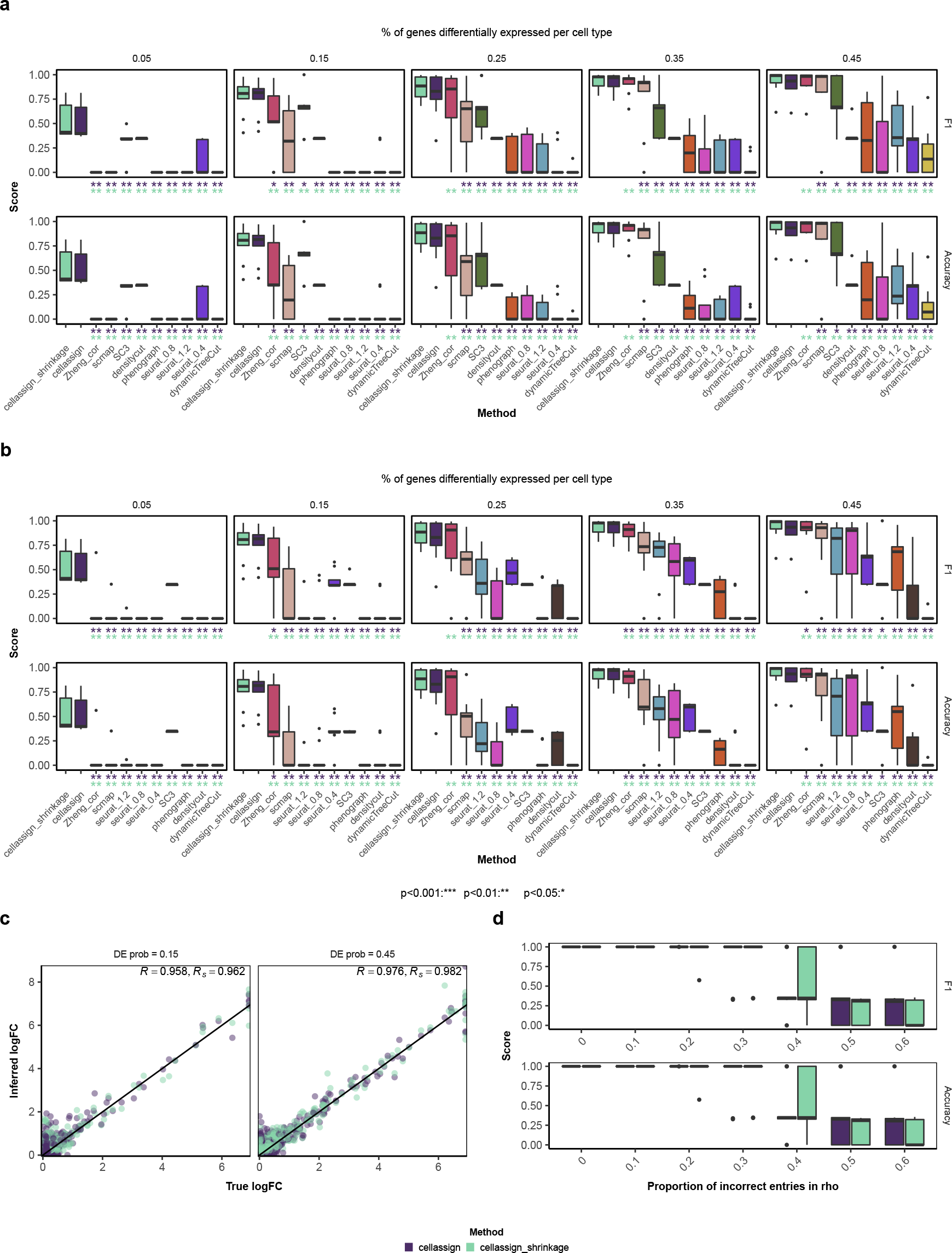
Performance of CellAssign on simulated data. (**a**) Accuracy and cell-level F1 score (**Methods**) for varying proportions of differentially expressed genes per cell type, with other differential expression parameters set to MAP estimates determined from comparing naïve CD8+ and naïve CD4+ T cells (**Methods**). CellAssign was provided with a set of marker genes (**Methods**); all other methods were provided with all genes. cellassign_shrinkage refers to a version of CellAssign with a shrinkage prior on *δ* (**Methods**). Asterisks indicate FDR-adjusted statistical significance (Wilcoxon signed-rank test) for pairwise comparisons between cellassign/cellassign_shrinkage and other methods. (**b**) Accuracy and cell-level F1 score for varying proportions of differentially expressed genes per cell type, with other differential expression parameters set to MAP estimates determined from comparing naïve CD8+ and naïve CD4+ T cells. All methods were provided with the same set of marker genes. (**c**) Correspondence between true simulated log fold change values and log fold change (*δ*) values inferred by CellAssign. *R* and *R*_*s*_ refer to the Pearson correlation between true and inferred logFC values for cellassign and cellassign_shrinkage, respectively. (**d**) Performance of CellAssign where a certain proportion of entries in the marker gene matrix are flipped at random. Differential expression parameters used for these simulations were based on those determined from comparing B and CD8+ T cells.

We next assessed the robustness of CellAssign to misspecification of marker gene information, acknowledging the likely scenario where user-provided marker gene information may be incomplete or incorrect. For example, a shared marker gene may be incorrectly specified as a cell type-specific marker gene due to incomplete prior information. We randomly flipped a proportion of entries in the binary marker gene matrix to introduce error. When supplied with data for 5 marker genes per cell type, CellAssign maintained comparable performance in scenarios where up to 30% of matrix entries were misspecified (**Figure 2D**, **Supplemental Table 1**). This robustness was maintained even when cells belonged to transcriptionally similar cell types containing fewer highly differentially expressed genes. For example, when cells were simulated based on the degree of dissimilarity between naïve CD4+ and naïve CD8+ T cells, CellAssign prediction accuracy was maintained in scenarios where 30% of marker gene matrix entries were misspecified (**Supplemental Figure 2A,B**, **Supplemental Table 1**).

We then evaluated performance of CellAssign on real scRNA-seq data from experimentally sorted populations. We applied CellAssign to data for FACS-purified H7 human embryonic stem cells in various stages of differentiation (8 cell types) [12]. Using bulk RNA-seq data from the same cell types, we defined a set of 84 marker genes for CellAssign based on differential expression results (**Supplemental Table 2**; **Methods**). CellAssign outperformed the most competitive unsupervised methods from systematic analysis (SC3, Seurat) [6] according to accuracy and cell type-level F1 score (**Supplemental Figure 3A-D,F**; **Methods**), with similar results obtained using only marker gene expression data (**Supplemental Figure 3E,G**) (CellAssign F1: 0.947, accuracy: 0.948; best unsupervised F1: 0.841, accuracy: 0.93). As an example of CellAssign’s ability to discriminate highly related cell types, anterior primitive streak (APS) and mid primitive streak (MPS) cells were accurately classified (83/84 correct), while no other method could reliably do so (all other methods assigned APS and MPS cells to the same cluster, **Supplemental Figure 3**).

**Figure 3.**
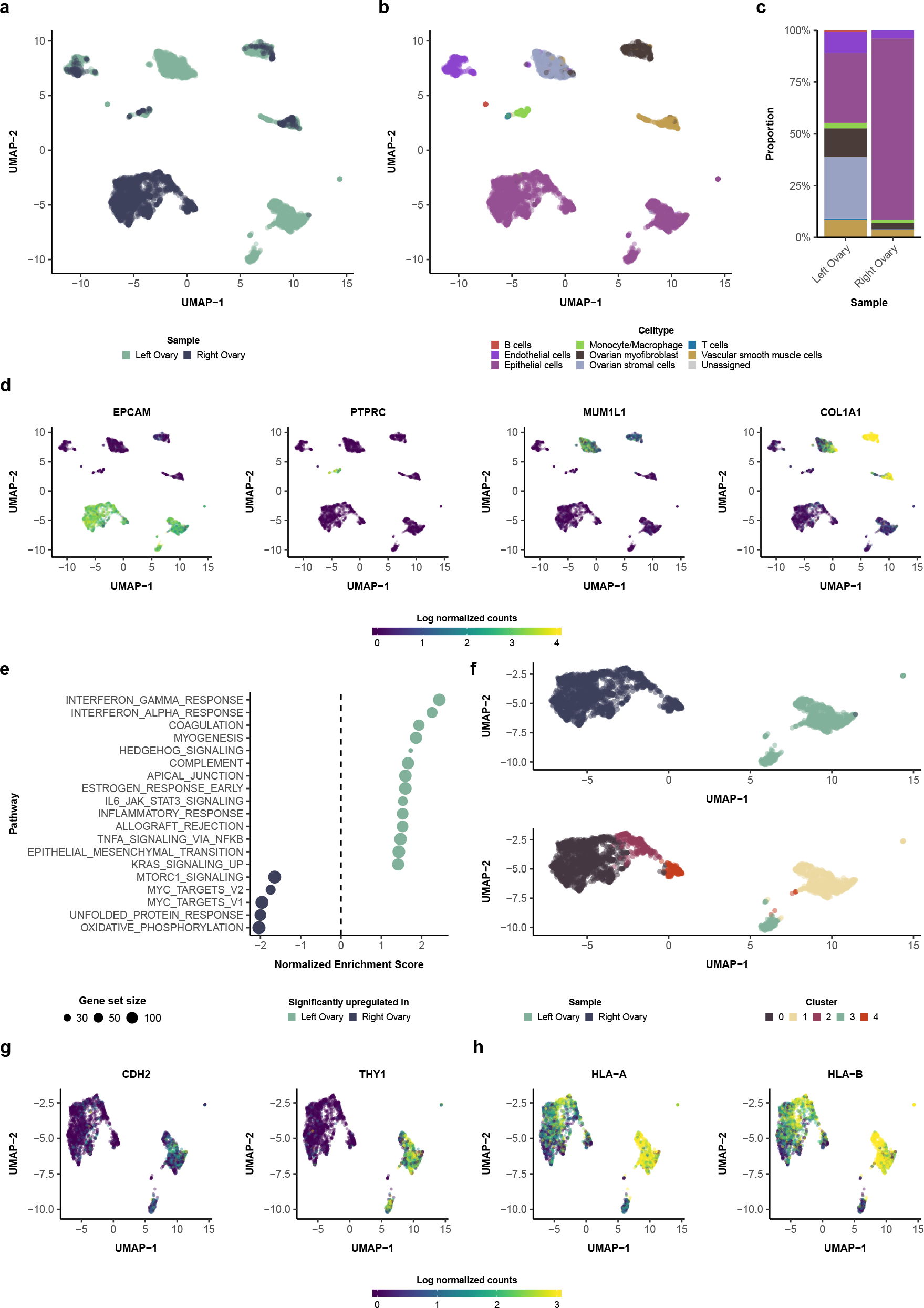
CellAssign infers the composition of the HGSC microenvironment. (**a**) UMAP plot of HGSC single cell expression data, labeled by sample. (**b**) UMAP plot of HGSC single cell expression data, labeled by maximum probability assignments from CellAssign. (**c**) Proportions of CellAssign cell types in each sample. (**d**) Expression (log normalized counts) of *EPCAM* (for epithelial cells), *CD45* (*PTPRC*) (for hematopoietic cells), *MUM1L1* (for ovary-derived cells), and *COL1A1* (for collagen-producing fibroblasts and smooth muscle cells). Expression values were winsorized between 0 and 4. (**e**) Hallmark pathway enrichment results for left ovary vs. right ovary epithelial cells (**Methods**). (**f**) Unsupervised clustering of epithelial cells (**Methods**). (**g**) Expression (log normalized counts) of epithelial-mesenchymal transition (EMT) associated markers, N-cadherin (*CDH2*) and *CD90* (*THY1*) in epithelial cells. (**h**) Expression (log normalized counts) of select HLA class I genes in epithelial cells.

### 2.3 Profiling the tumour microenvironment composition of spatially sampled HGSC

We next exemplified CellAssign in a real world setting where the aim was to decompose cancer tissues from patients into constituent microenvironmental components and profile variation across anatomic space and between malignant clones. We generated scRNA-seq data for 5233 cells from 2 spatial sites from an untreated high-grade serous ovarian cancer patient at the time of primary debulking surgery (left ovary: 2818 cells, right ovary: 2415 cells). From this data, we executed an analytic workflow consisting of dimensionality reduction, assignment of cell types and differential expression between malignant cell subpopulations.

We began analysis by dimensionality reduction with uniform manifold approximation and projection (UMAP [17]) revealing four major site-specific populations and four mixed populations with representation from both samples (**Figure 3A**). Using a panel of literature-derived marker genes (**Supplemental Table 2**, **Methods**), we identified 8 major epithelial, stromal, and immune cell types with CellAssign (**Figure 3B,C**), which were consistent with well-known marker gene expression (**Figure 3D**, **Supplemental Figure 4A**, **Methods**). Ovarian stromal cells and myofibroblasts were identified based on expression of *MUM1L1*, *ARX*, and *KLHDC8A*, ovary-specific markers known to be expressed in stroma from bulk RNA-seq and immunohistochemistry [18] (**Figure 3D**, **Supplemental Figure 4A**), with myofibroblasts distinguished by higher expression of *α*-smooth muscle actin and various collagen genes [19] (**Figure 3D**, **Supplemental Figure 4A**). Unlike other non-epithelial cell types, ovarian stromal cells were largely restricted to the left ovary. A group of cells expressing vascular smooth muscle markers *α*-smooth muscle actin, *MYH11*, and *MCAM* [20] was also identified with CellAssign (**Supplemental Figure 4A**). We note that for cell types such as ovarian stromal cells, no scRNA-seq data from purified populations was available. Thus, CellAssign can annotate TME cell types for which marker genes have been orthogonally derived in the literature but scRNA-seq data for purified populations is unavailable. Hematopoietic cells (B cells, T cells, and myeloid cells) were rare in both samples (left ovary: 4%, right ovary: 1.5%; **Figure 3C**) and dominated by myeloid populations (65.7% and 87.9% of hematopoietic cells in left and right ovary, respectively). While CellAssign resolved hematopoietic cell types in a manner consistent with the expression patterns of canonical marker genes, unsupervised approaches did not resolve some of these cell types, such as B cells, from other hematopoietic or non-hematopoietic cell types (**Supplemental Figure 5**). Thus for TME decomposition and profiling, subtle differences between constituent cell types may be better distinguished by CellAssign over standard approaches [21].

We next characterized variation within the epithelial cells determined by CellAssign, all of which were determined to be malignant based on ubiquitous expression of epithelial ovarian cancer markers [22, 23] (**Supplemental Figure 4B**). Within epithelial cells we identified five clusters using Seurat (**Figure 3F**) with three (0, 2, 4) derived from the right ovary and two (1, 3) from the left ovary. Differential expression between clusters revealed significant upregulation of genes associated with epithelial-mesenchymal transition in the left ovary (*Q* = 0.021), including N-cadherin (*CDH2*) and *CD90* (*THY1*) (**Figure 3E-G**), and downregulation of E-cadherin (*CDH1*; *Q* = 4.8e-19). Immune-associated pathways were also significantly upregulated, primarily due to cluster 1, one of the two clusters from the left ovary (**Figure 3E,F,H**, **Supplemental Figure 6A**, **Supplemental Figure 7A-B**, **Methods**). HLA class I genes were amongst the most differentially expressed genes associated with these pathways (**Supplemental Figure 7B**). While HLA expression in cluster 1 was comparable to levels in stromal cells and myofibroblasts, expression levels in other clusters were lowest across all cell types (**Supplemental Figure 6B**), suggestive of subclonal HLA downregulation. We next considered cluster-specific gene expression among epithelial cells in the right ovary. Hypoxia response was significantly upregulated in cluster 2 relative to the other right ovary clusters (all *Q* < 7e-04; **Supplemental Figure 7C-E**). Accordingly, apoptosis and glycolysis pathways were also upregulated while cell cycle and oxidative phosphorylation-associated pathways were downregulated, consistent with hypoxia-induced cell cycle arrest and metabolic dependence on glycolysis (**Supplemental Figure 7C,D**). Together, TME and malignant cell profiling of multi-site HGSC samples demonstrate on real data how CellAssign can be leveraged within analytical workflows, superseding standard clustering approaches to decompose the TME without compromising the ability to characterize variation within major cell types.

### 2.4 Temporal immune microenvironment dynamics accompanying follicular lymphoma progression and transformation

To dissect microenvironmental changes in follicular lymphoma tracking with disease progression and transformation, we sequenced the transcriptomes of 9754 cells from temporally collected lymph node biopsies of 2 follicular lymphoma patients at two time points each (4 samples total). Histopathological transformation to diffuse large B cell lymphoma (DL-BCL) occurred in one patient (FL1018), while progression occurred in the other (FL2001) 2 years after rituximab treatment (4 years after diagnosis; **Figure 4A**). We then investigated temporal phenotypic dynamics using CellAssign to establish cell type composition of malignant and nonmalignant cells in the microenvironment.

**Figure 4.**
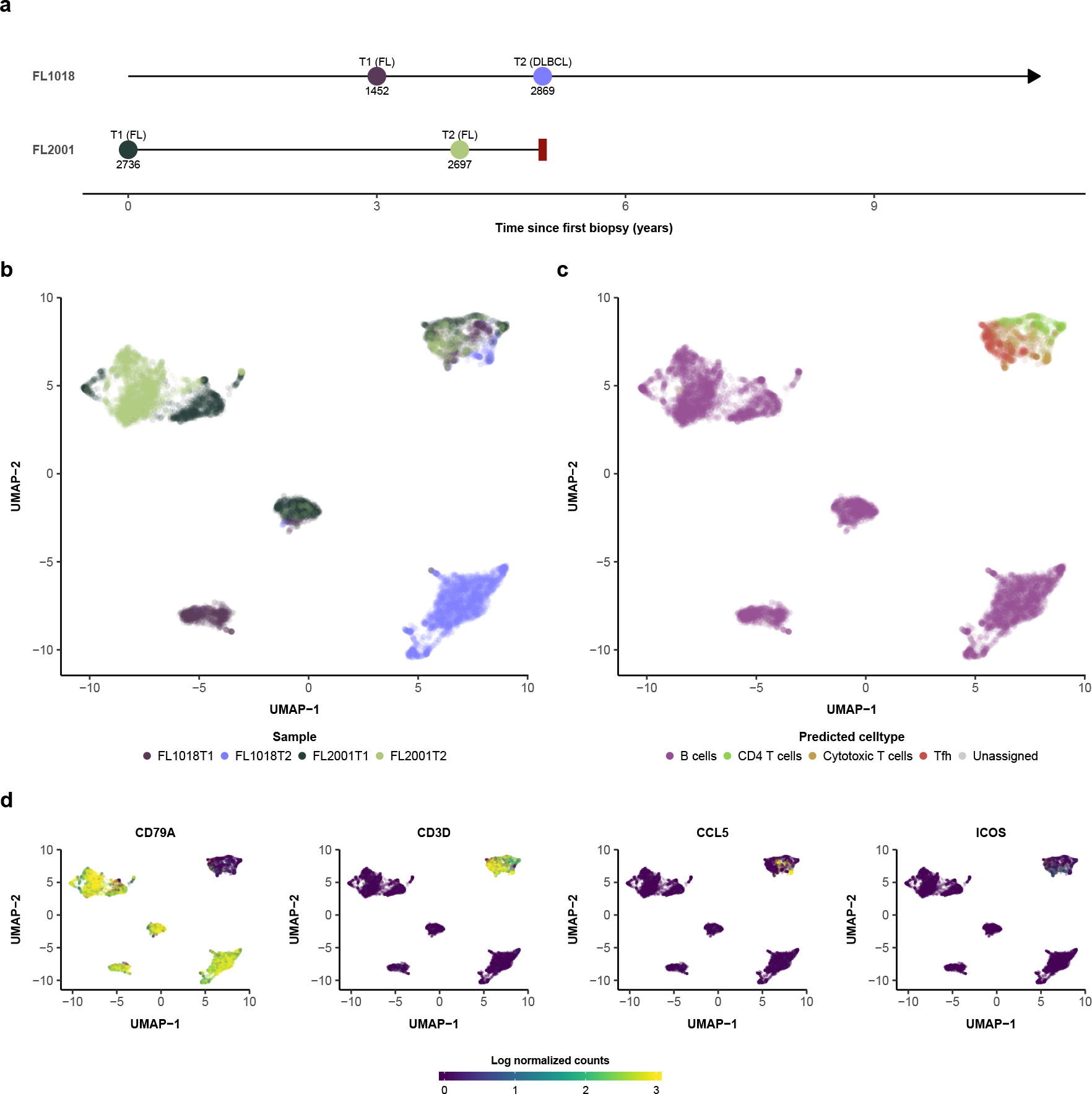
CellAssign infers the composition of the follicular lymphoma microenvironment.(**a**) Sample collection times for FL1018 (transformed FL) and FL2001 (progressed FL). FL1018 is alive while FL2001 was lost to followup (indicated by the red rectangle). The number of cells collected for each sample is indicated. (**b**) UMAP plot of follicular lymphoma single cell expression data, labeled by sample. (**c**) UMAP plot of follicular lymphoma single cell expression data, labeled by maximum probability assignments from CellAssign. (**d**) Expression (log normalized counts) of select marker genes *CD79A* (for B cells), *CD3D* (for T cells), *CCL5* (for CD8+ T cells), and *ICOS* (for T follicular helper cells). Expression values were winsorized between 0 and 3.

We first applied UMAP which yielded three major patient-specific and two mixed populations comprised of cells from both patients (**Figure 4B**). Leveraging literature-derived marker gene information (**Supplemental Table 2**), we applied CellAssign to identify 4 major T and B cell types (**Figure 4C,D**, **Supplemental Figure 8**, **Methods**). In comparison, most unsupervised approaches were unable to cleanly resolve T cell subpopulations in the microenvironment (**Supplemental Figure 9**), thereby reducing the capacity to interpret a crucial component of the lymphoma TME. We surmised the mixed B cell population likely contained nonmalignant B cells (**Figure 5A**), and accordingly we examined immunoglobulin light chain constant domain expression using CellAssign (**Figure 5B**) to confirm heterogeneous light chain expression (*κ*/*IGKC* or *λ*/*IGLC*) in the polyclonal nonmalignant B cell population and homogeneous light chain restriction in the clonally identical malignant B cell population (‘light chain restriction’) [24]. The three patient-specific B cell populations were largely IGLC positive, consistent with malignant expansion of *λ*-chain expressing cells. Applying CellAssign to the mixed population (**Supplemental Table 2**) showed that 456/774 cells (58.9%) were IGKC+ (FL1018: 67/106 (63.2%), FL2001: 389/668 (58.2%)), consistent with the expected polyclonal 60:40 ratio in normal lymphoid organs [25] (**Supplemental Figure 10**). In addition, scRNA-seq data of reactive lymph node (RLN) B cells from four healthy donors mapped onto the mixed B cell population [26], (**Figure 5C**, **Supplemental Figure 11**). This population also expressed significantly lower levels of follicular lymphoma markers *BCL2* and *BCL6* [27–28, 29, 24] than the other B cells (all *Q* < 1.8e-07; **Supplemental Figure 12**, **Supplemental Table 3**). Together these results demonstrate the ability of CellAssign to distinguish malignant from nonmalignant B cells, thereby enhancing cell decomposition capacity and cell type interpretation for lymphoid cancers.

**Figure 5.**
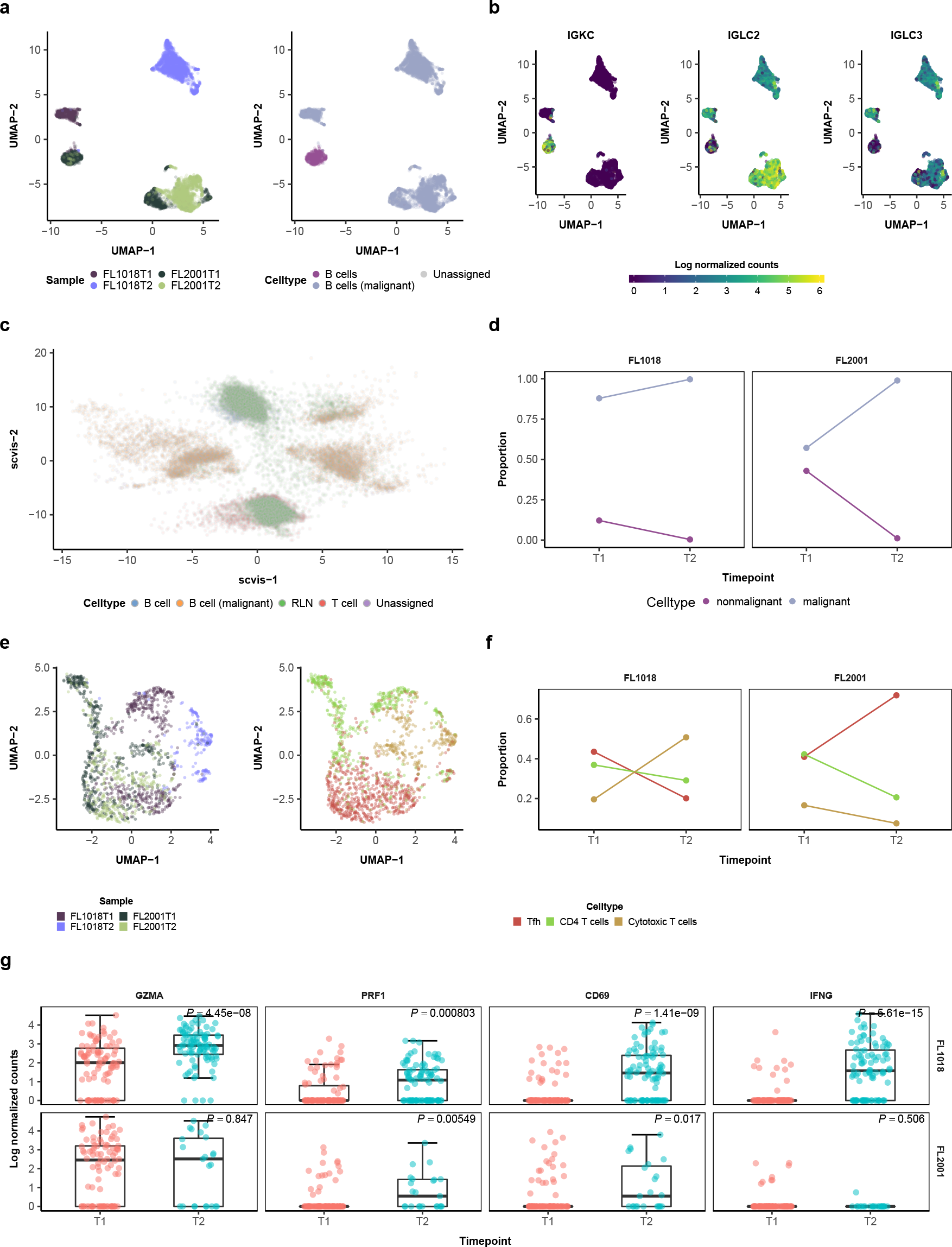
Temporal changes in nonmalignant cells in the follicular lymphoma microenvironment. (**a**) Left: UMAP plot of CellAssign-inferred B cells, labeled by sample. Right: UMAP pot of CellAssign-inferred B cells, labeled by putative malignant/nonmalignant status. (**b**) Expression (log normalized counts) of *κ* (*IGKC*) and *λ* (*IGLC2* and *IGLC3*) light chain constant region genes. Expression values were winsorized between 0 and 6. (**c**) Scvis plot of follicular lymphoma data and single cell RNA-seq data of lymphocytes from reactive lymph nodes from healthy patients. The follicular lymphoma data was used to train the variational encoder and produce the two-dimensional embedding. Indicated cell types are B cell (nonmalignant B cell from FL), B cell (malignant) (malignant B cell from FL), T cell (T cell from FL), RLN (reactive lymph node cell). (**d**) Relative proportion of B cell subpopulations over time. (**e**) UMAP plots of FL T cells, labeled by sample and CellAssign-inferred celltype. (**f**) Relative proportion of T cell subpopulations over time. (**g**) Normalized expression of CD8+ T cell activation markers over time. *P* -values computed with the Wilcoxon rank-sum test and adjusted with the Benjamini-Hochberg method.

We next investigated the temporal dynamics of these cell types in the two patients. The relative proportion of nonmalignant B cells decreased dramatically over time in both cases (FL1018: 12.2% to 1.4%; FL2001: 44.4% to 1.4%) (**Figure 5D**), consistent with clonal expansion of malignant cells during disease progression. Among T cells, the relative proportions of each cell type were comparable between patients in diagnostic samples, but exhibited divergent trajectories following recurrence, with cytotoxic T cells dominating the transformed sample while T follicular helper cells dominated the progressed sample (**Figure 5E,F**).

We examined whether these compositional changes were accompanied by cell type-specific phenotypic changes. Differential expression analysis revealed significant upregulation of immune-associated pathways such as cytokine signalling [30] and T-cell activation and effector molecules among cytotoxic T cells, T follicular helper cells, and CD4+ T cells after transformation (*CD69* in all T cells, *IFNG*, *GZMA*, and *PRF1* in cytotoxic T cells [31]; **Supplemental Figure 13A**, **Figure 5G**, **Supplemental Figure 14**, **Supplemental Table 3**). Thus, transformation in this case appeared to be accompanied by T-cell activation.

Within malignant cells, upregulation of cell cycle-associated pathways (E2F targets, and G2M checkpoint; all *Q* < 0.0016) and an increase in the proportion of cycling (S or G2/M phase) malignant cells was observed upon transformation, suggesting an increase in replicative potential [32] (**Figure 6A,C,D**, **Supplemental Table 3**). Several immune-associated pathways, including complement and interferon gamma response, were significantly downregulated upon transformation (**Figure 6A**, **Supplemental Table 3**). To interpret these findings, we enumerated genes most downregulated upon transformation based on log-fold change and significance (**Supplemental Figure 15**), which included several HLA class I and II genes (**Figure 6G**). While HLA expression levels in nonmalignant B cells were similar between timepoints (**Figure 6G,H**), malignant cells had significantly lower HLA expression at transformation (all *Q* < 9.6e-24), and the HLA class I antigen presentation pathway was downregulated upon transformation in malignant cells (*Q* = 0.019; **Supplemental Figure 16**). Coupled with the increase in cytotoxic T cell proportion and upregulation of T-cell activation markers upon transformation, these results are consistent with immune escape following transformation.

**Figure 6.**
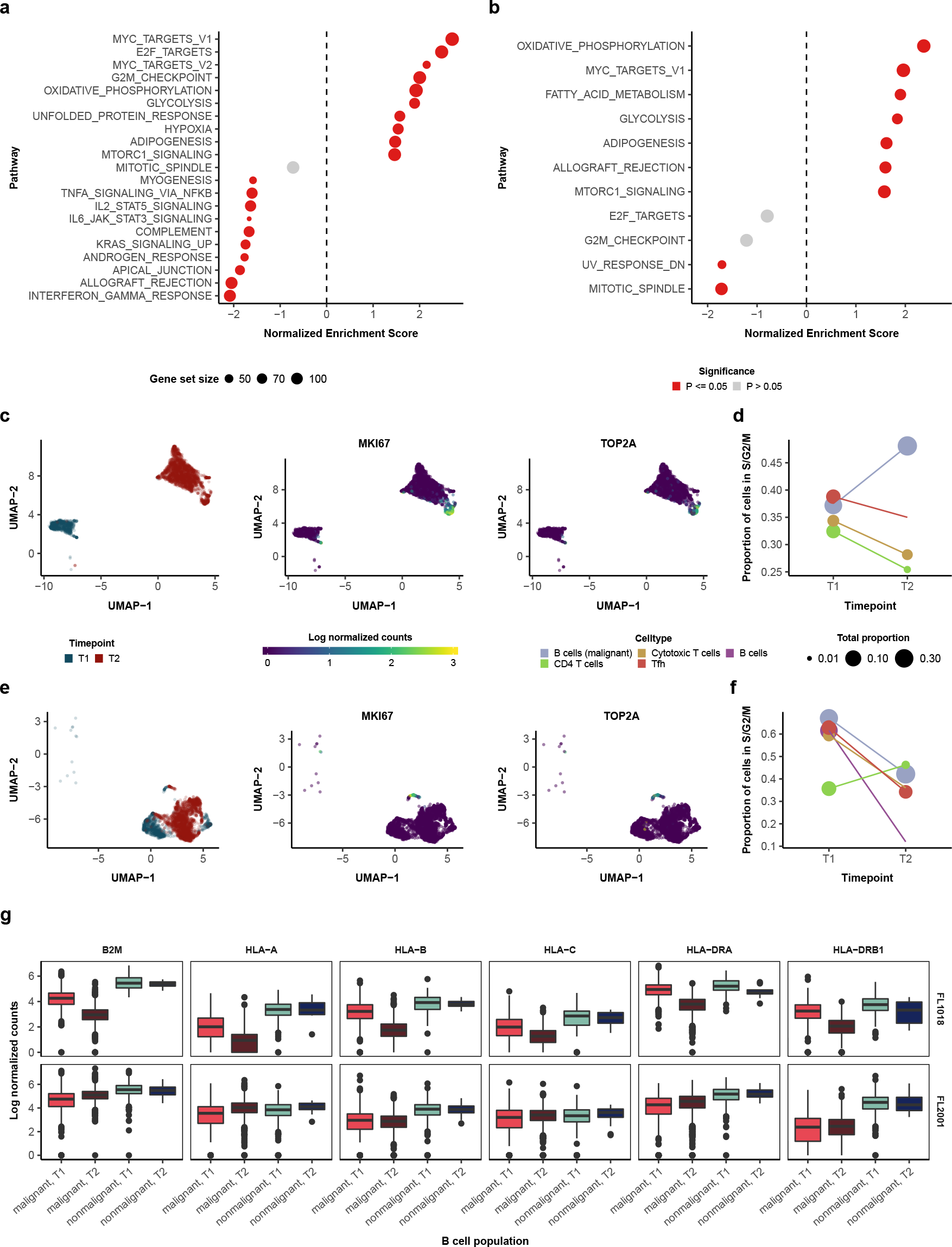
Temporal changes in malignant cells in the follicular lymphoma microenvironment. (**a,b**) Pathway enrichment scores computed by fGSEA for differentially enriched (adjusted *P* ≤ 0.05) and cell cycle-associated pathways among malignant cells between timepoints for (a) FL1018 and (b) FL2001 (**Methods**). Pathways with a positive enrichment score are upregulated in T2 compared to T1 samples. *P* -values were adjusted with the Benjamini-Hochberg method. (**c,e**) UMAP plots, labeled by sample and proliferation marker expression (*MKI67* and *TOP2A*), for (c) FL1018 and (e) FL2001. Expression values were winsorized between 0 and 3. (**d,f**) Proportion of cells assigned to be in non-G1 cell cycle phases (S/G2/M) by cyclone across timepoints in (d) FL1018 and (f) FL2001. Normalized expression of HLA class I genes and select HLA class II genes across timepoints in FL1018 and FL2001.

In FL2001 (progressed FL), fewer pathways were differentially expressed between timepoints in malignant cells, indicative of more stable dynamics (**Figure 6B**). The cell cycle-associated mitotic spindle pathway was downregulated upon early progression (*Q* = 0.0043; **Figure 6B**). Concordantly, cell cycle analysis [33] revealed an decrease in the proportion of cycling (S or G2/M phase) malignant cells after progression (**Figure 6E,F**). Together, these results illustrate how CellAssign can be feasibly applied to study compositional and phenotypic changes in the tumour microenvironment at the level of individual cell types, and the clonal dynamics of malignant populations using non-canonical marker genes tailored to identifying specific populations of interest.

## 3 Discussion

We developed a computational method to automatically annotate single cell RNA sequencing data into cell types based on pre-defined marker gene information. Our approach systematically determines cell type expression patterns and assignment probabilities based solely on the assumption that marker genes are highly expressed in their respective cell types, eliminating the need for manual cluster annotation or existing training data for cell type mapping methods. In simulations and on real scRNA-seq data from purified populations, CellAssign’s accuracy was comparable or superior to state-of-the-art workflows based on unsupervised clustering and mapping methods, and ran in a minute on datasets of tens of thousands of cells. We additionally demonstrate how bulk RNA-seq data can enable marker gene identification for accurate discrimination of phenotypically similar cell types with CellAssign.

We subsequently applied CellAssign to dissect the microenvironment composition of spatially- and temporally-collected samples from HGSC and follicular lymphoma. We show how CellAssign can not only delineate multiple malignant and nonmalignant epithelial, stromal, and immune cell types, but also identify subpopulations defined by arbitrary marker genes, uncovering *IGKC*:*IGLC* ratios among nonmalignant B cells in follicular lymphoma consistent with those for normal lymphoid structures [25]. While these analyses are constrained by restricted cohort size, they provide first-of-kind examples of spatiotemporal dynamics and microenvironment interplay interpreted through leveraging prior knowledge of cell types in a prinicipled statistical approach.

We note that CellAssign is intended for scenarios where well understood marker genes exist. Poorly characterized cell types (or unknown cell types or cell states) may be invisible to the CellAssign approach. Furthermore, we make no *a priori* distinction between “medium” or “high” expression of the same marker in two different cell types, though these could be incorporated by extending the model to accommodate constraints between different *δ* parameters. Nevertheless, we suggest a large proportion of clinical applications profiling complex tissues start with hypotheses relating the composition of known cell types to disease states. As such, CellAssign fills an important role in the scRNA-seq analysis toolbox, providing interpretable output from biologically motivated prior knowledge that is immune to common issues plaguing unsupervised clustering approaches [8].

The volume of scRNA-seq data will increase over time in two important ways: (i) the number of cell types profiled will increase, thereby expanding databases of known marker genes and (ii) scRNA-seq data will become more widely available in research and clinical settings [34]. CellAssign is therefore poised to provide scalable, systematic and automated classification of cells based on known parameters of interest, such as cell type, clone-specific markers, or genes associated with drug response. Furthermore, by appropriately modifying the observation model CellAssign can easily be extended to annotate cell types in data generated by other single-cell measurement technologies such as mass cytometry. We anticipate the CellAssign approach will help unlock the potential for large scale population-wide studies of cell composition of human disease and other complex tissues through encoding biological prior knowledge in a robust probabilistic framework.

## 4 Acknowledgements

We thank Dr. Valentine Svensson (Caltech) for his feedback on this manuscript. We also thank Dr. Wyeth Wasserman (BC Children’s Hospital), Dr. Brad Nelson (Deeley Research Centre), Dr. Phineas Hamilton (Deeley Research Centre), and Dr. Alex Miranda Rodriguez (Deeley Research Centre) for helpful discussions.

## 5 Funding

A.W.Z. is funded by scholarships from the Canadian Institutes of Health Research (CIHR) (Vanier Canada Graduate Scholarship, Michael Smith Foreign Study Supplement) and a BC Children’s Hospital-UBC MD/PhD Studentship. K.R.C. is funded by postdoctoral fellowships from the CIHR (Banting), the Canadian Statistical Sciences Institute (CANSSI), and the UBC Data Science Institute. S.P.S. is a Susan G. Komen scholar. We acknowledge generous funding support provided by the BC Cancer Foundation. In addition, S.P.S. receives operating funds from the CIHR (grant FDN-143246), Terry Fox Research Institute (grants 1021 and 1061) and the Canadian Cancer Society (grant 705636). This work was supported by Cancer Research UK grant C31893/A25050 (S.A. and S.P.S.). S.P.S is supported by the Nicholls-Biondi endowed chair and the Cycle for Survival benefitting Memorial Sloan Kettering Cancer Center. CS is an Allen Distinguished Investigator supported by the Allen Frontiers Group.

## 6 Author Contributions

Study design: A.W.Z., K.R.C., S.P.S.; Writing: A.W.Z., K.R.C., S.P.S.; Manuscript review: A.W.Z., C.O., E.A.C., J.L.P.L., A.M., M.W., T.A., A.P.W., J.N.M., S.A., C.S., K.R.C., S.P.S.; Data interpretation: A.W.Z., S.A., C.S., K.R.C., S.P.S.; Data curation: B.H., D.L., L.C.; Data analysis: A.W.Z., K.R.C., M.W., P.W., T.C., X.W.; Model development: A.W.Z., K.R.C., S.P.S.; Single cell processing: C.O., E.A.C., J.L.P.L.; Case identification: A.M., J.N.M., C.S.; Supervision: K.R.C., S.P.S., C.S., S.A.

## 7 Competing Interests

S.P.S. and S.A. are founders, shareholders, and consultants of Contextual Genomics Inc.

## 8 Methods

### 8.1 Ethics

Ethical approval for this study was obtained from the University of British Columbia (UBC) Research Ethics Board (ethics numbers H08-01411, H14-02304, and H18-01090).

### 8.2 The CellAssign model

#### 8.2.1 Model description

Let **Y** be a cell-by-gene expression matrix of raw counts for *N* cells and *G* genes. Suppose among those cells we have *C* total cell types, each of which is defined by high expression of several “marker” genes. We encode the relationship between cells and marker genes through a binary matrix ***ρ***, where *ρ*_*gc*_ = 1 if gene *g* is a marker for cell type *c* and 0 otherwise. To relate cells to cell types, we introduce an indicator vector **z** = {*z*_*n*_} that encodes to which of the *C* cell types each cell belongs:

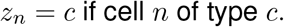

In order to assign cells to cell types we perform statistical inference of the probability that each cell is of a given cell type for which we must compute the quantity 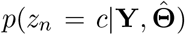, where 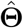 are the MAP estimates of the model parameters.

Let *s*_*n*_ be the size factor for cell *n* and **X** be a *P × N* matrix of *P* covariates (such as patient of origin). then our model is

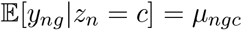

where

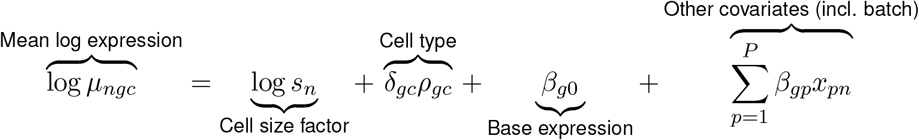

with the constraint that *δ*_*gc*_ > 0.

The intuition here is that if gene *g* is a marker for cell type *c* then we expect the expression of *g* to be multiplied by the factor 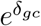, where *δ*_*gc*_ is inferred. In this way we put no restriction that marker genes can’t be expressed in other cell types and that they must be highly expressed in their cell type, only that they exhibit higher expression in the cells of type for which they are a marker. The quantity *δ*_*gc*_ corresponds to the average log fold change that gene *g* is over-expressed in cell *c*, which only occurs for marker genes for cell types since *ρ*_*gc*_ must equal 1 for this to contribute to the likelihood. By default we impose a lower bound such that *δ* > log 2, making the interpretation that a marker gene must be over-expressed by a factor of 2 relative to cells for which it is not a marker, but this is left as an option for the user. We also control for technical or sample effects through the matrix **X**.

The user can specify whether or not to put a lognormal shrinkage prior over *δ*_*gc*_ values 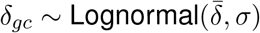, where the mean and variance parameters of the lognormal 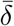 and *σ* are initialized to 0 and 1, respectively. In plot labels, `cellassign_shrinkage` refers to the version of CellAssign with this option turned on.

#### 8.2.2 Inference

The likelihood is given by

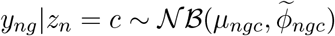

where 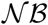 is the negative binomial distribution parametrized by a mean *μ* and a *μ*-specific dispersion 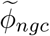. We define 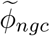 as a sum of radial basis functions dependent on the modelled mean *μ*_*ngc*_ as proposed by a recent publication [35]:

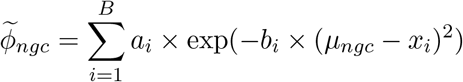

where *a*_*i*_ and *b*_*i*_ represent RBF parameters to be fitted, *B* is the total number of *centers* in the RBF, and *x*_*i*_ is center *i*. The centers are set to be equally spaced apart from 0 to the maximum number of counts max *y*_*ng*_.

Using EM for inference, the latent variables are **z** ≡ {*z*_*n*_} while the model parameters to be maximized are ***δ*** = {*δ*_*gc*_}, ***β*** = {*β*_*g*0_, *β*_*gp*_}, **a** = {*a*_*i*_}, and **b** = {*b*_*i*_}.

##### E-step

Compute

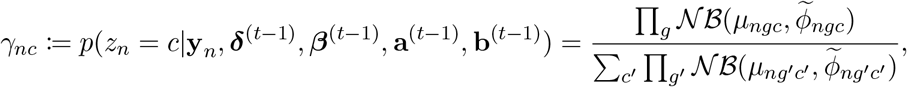

where ***θ***^(*t*)^ is the value of some parameter ***θ*** at iteration *t*. We then form the *Q* function

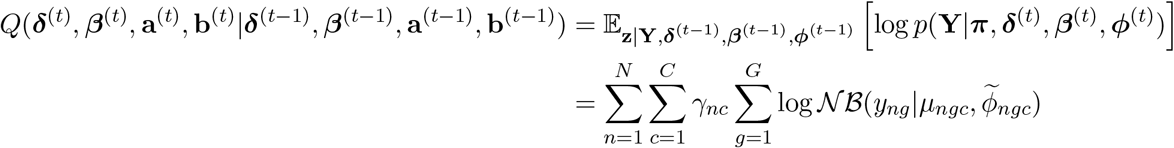

##### M-step

During the M-step we optimize the above *Q*-function using the ADAM optimizer as implemented in Google’s Tensorflow [37]. By default we use a learning rate of 0.1, allow a maximum of 10^5^ ADAM iterations per M-step, and consider the M-step converged when the relative change in the *Q* function value falls below 10^−4^. By default we consider the EM algorithm converged when the relative change in the marginal log likelihood falls below 10^−4^.

##### Initialization

The following initializations are used for model parameters:

- *β*_*gp*_ is drawn from a 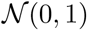 distribution
- log *δ*_*gc*_ is drawn from a 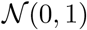 distribution truncated at [log(*δ*_min_), 2]
- *a* is initialized to 0
- *b* is initialized to twice the square difference between successive spline bases

#### 8.2.3 Availability

CellAssign is available as an R package at github.com/irrationone/cellassign.

### 8.3 Simulation

#### 8.3.1 Model description and rationale

Initially, we attempted to simulate multi-group data from the `splatter` model. We employed 10x Chromium data for peripheral blood mononuclear cells (PBMC) [16] with cell type labels derived from [38] to determine realistic parameter estimates for the differential expression component of the model (see below). In order to do so, group-specific log fold-change (logFC) values were drawn from a mixture distribution of a central, narrow Gaussian-Laplace mixture (representing non-differentially expressed genes) and two flanking, absolute value-transformed Gaussians (representing downregulated/upregulated genes). This mixture distribution was fitted to logFC values derived from differential expression analysis (see below).

However, inspection of posterior predictive samples for multiple fits, using labeled singlecell RNA-seq data from [16] and FACS-purified data from Koh et al. [12] (**Supplemental Figure 17A,B**, **Supplemental Figure 18A,B**), revealed that this model systematically underestimates extreme logFC values (**Supplemental Figure 17C**, **Supplemental Figure 18C**). Thus, to accommodate the heavier tails present in observed data, we augmented the splatter model by replacing the flanking absolute value-transformed Gaussian components with bounded Student’s *t* distributions. Posterior predictive logFC distributions from this modified model better fit the observed data (**Supplemental Figure 17D**, **Supplemental Figure 18D**). Consequently, we used this model to perform simulation analysis.

#### 8.3.2 Model fitting

The models described above were fit to logFC values derived from real data. Using the labeled 10x Chromium data for 68k PBMCs [16], differential expression was performed with the `findMarkers` function from the R package scran [39]. To generate corresponding null distributions of logFC values for non-differentially expressed genes, we split data for each cell type into equally sized halves 10 times, running findMarkers to compare the resulting halves. A central Gaussian-Laplace mixture (*μ* = 0) was first fit to the null logFC values. The distribution of posterior predictive logFC values appeared to be consistent with observed logFC values for this null component (**Supplemental Figure 17D**). Following this, the entire mixture distribution was fitted to logFC values for pairs of distinct cell types, using *maximum a posteriori* (MAP) estimates of parameters for the central Gaussian-Laplace component. Posterior distributions of model parameters were inferred using the no U-turn sampler (NUTS) in pymc3, using 4 independent chains, 1000 tuning iterations, and 2500 additional iterations per chain. Trace plots and the Gelman-Rubin diagnostic were used to assess convergence.

#### 8.3.3 Simulating multi-group data

Expression count matrices were simulated using a modified version of the `splatter` package. Log fold change values were simulated according to our model instead of the splatter model. Other settings were kept identical. We used MAP estimates of *μ*_+_, *μ_−_*, *σ*_+_, *σ_−_*, *ν*_+_, and *ν_−_*, determined by fitting our simulation model to (1) logFC values between naïve CD4+ and naïve CD8+ T cells (**Supplemental Figure 17A**); and (2) logFC values between B cells and CD8+ T cells (section 8.3.1) for the differential expression component. The proportion of downregulated genes out of differentially expressed genes was set to 0.5 (i.e. equally probable for a differentially expressed gene to be downregulated vs. upregulated). Three “groups” (cell types) were simulated at equal proportions. Other parameters for splatter were fitted from 10x Chromium data for 4,000 T cells available from 10x Genomics.

To assess the performance of CellAssign relative to other clustering methods across a range of *p*_*d*_ values (proportion of genes differentially expressed between each pair of cell types), *p*_*d*_ was chosen from {0.05, 0.15, 0.25, 0.35, 0.45, 0.55}. (The true MAP estimate of *p*_*d*_ was 0.0746 for naïve CD4+ vs. naïve CD8+ T cells, and 0.153 for B vs. CD8+ T cells.) The number of simulated cells, *n*, was set to 2000, and 1000 were randomly set aside for training (for scmap and correlation-based supervised clustering).

To assess the robustness of CellAssign to misspecification of the marker gene matrix *ρ*, *p*_*d*_ was set to 0.25 and the number of simulated cells *n* to 1500.

Simulations were run 9 times with unique random seeds for each combination of parameter settings.

#### 8.3.4 Clustering multi-group data

Count matrices were normalized with scater normalize and the top 50 principal components were computed from the top 1000 most variable genes. For `phenograph, Seurat` (resolution ∊ {0.4, 0.8, 1.2}), `densitycut`, and `dynamicTreeCut`, unsupervised clustering was performed on the values of these top 50 PCs. For `SC3`, the entire normalized SingleCellExperiment object was passed as input instead. For supervised methods (`scmap-cluster` [10] and correlation-based [16]), expression data for both training and evaluation sets was provided. For CellAssign, the raw count matrix was provided as input, along with a set of marker genes selected based on simulated log fold change and mean expression values. Specifically, a gene was defined as a marker gene if it was in the top 5th percentile of differentially expressed genes according to logFC and the top 10th percentile of differentially expressed genes according to mean expression. A maximum of 15 marker genes were selected for each group. In simulations of robustness to marker gene misspecification, a proportion of randomly selected entries in the marker gene matrix *ρ* were flipped from 0 to 1 (or vice versa). All other parameters were set to the defaults.

#### 8.3.5 Mapping clusters to true groups

For assignments derived from unsupervised clustering, clusters were mapped to simulated groups by first performing differential expression between each cluster and the remaining cells. Following this, we computed the Spearman correlation between these logFC values and the simulated (true) logFC values for each simulated group. Each inferred cluster was mapped to most highly correlated simulated group based on Spearman’s *ρ* where *ρ* > 0 and *P* ≤ 0.05. Clusters that could not be mapped based on these criteria were marked as ‘unassigned’.

#### 8.3.6 Evaluation

Accuracy and cell-level F1 score were computed to evaluate clustering performance. The cell-level F1 score considers each cell as an individual classification task with a true cell type assignment (and potentially multiple incorrect cell type assignments) for the purposes of calculating precision and recall.

#### 8.3.7 Benchmarking

We generated synthetic datasets for benchmarking from the modified splatter model (section 8.3.1) with Student’s *t* parameters *μ* = 0.1, *σ* = 0.1, *ν* = 1 and the proportion of differentially expressed genes per cell type set to 20%. Synthetic datasets of various sizes (number of cells *N* ∊ {1000, 2000, 4000, 8000, 10000, 20000, 40000, 80000} and number of cell types *C* ∊ {2, 4, 6, 8}) with a balanced number of cells per type were generated. Markers for CellAssign were selected from genes in the top 20th percentile in terms of log fold change among differentially upregulated genes and the top 10th percentile in terms of expression. CellAssign was run with 2, 4, 6, and 8 markers per cell type, with a maximum minibatch size of 5000 cells. On simulated data for 80000 cells from 2 cell types, CellAssign completed in under 2 minutes, appearing to scale at worst linearly in the number of cell types and marker genes used per cell type (**Supplemental Figure 19**). Five separate CellAssign runs were timed for each combination of parameters.

### 8.4 Koh *et al.* dataset

#### 8.4.1 Preprocessing and normalization of single-cell RNA-seq data

Preprocessed data was obtained from the R package `DuoClustering2018` [12, 6]. Cell-types with both single-cell RNA-seq data and bulk RNA-seq data were used: hESC (day 0 human embryonic stem cell), APS (day 1 anterior primitive streak), MPS (day 1 mid primitive streak), DLL1pPXM (day 2 DLL1+ paraxial mesoderm), ESMT (day 3 somite), Sclrtm (day 6 sclerotome), D5CntrlDrmmtm (day 5 dermomyotome), D2LtM (day 2 lateral mesoderm). Normalization and dimensionality reduction was performed with `scater normalize`, `runPCA`, `runTSNE`, and `runUMAP`. The top 500 most variable genes were used to compute the top 50 principal components, and the top 50 PCs were used as input for t-SNE.

#### 8.4.2 Identification of marker genes from bulk RNA-seq data

Differential expression analysis results for bulk RNA-seq data for the same cell types was used to compute the relative expression of each gene in each cell type. Briefly, bulk RNA-seq log fold change values obtained from [12] were used to compute log-scale relative gene expression levels. Next, we identified gene-specific thresholds for defining the cell types in which each gene is a marker. For each gene, relative expression levels across cell types were sorted in ascending order, denoted as *E*_1_,…, *E*_*C*_, where *C* is the total number of cell types. The maximum difference between sorted expression levels, max_1≤*i*<*C*_ (*E*_*i+1*_ − *E*_*i*_), was then computed. Denote the index *i* for gene *g* in which this difference is maximal *i*_*g*_. For gene *g*, cell types in which relative expression values were equal to or greater than 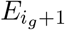 were considered cell types with gene *g* as a marker. Genes with a maximum difference value in the the top 20th percentile were used as marker genes.

#### 8.4.3 CellAssign

CellAssign was run on count data using the marker gene matrix defined from bulk RNA-seq data described above. Three random initializations of expectation-maximization were used with shrinkage priors on *δ*_*gc*_ turned on (section 8.2.1). Results from the run that reached the highest marginal log-likelihood at convergence were kept.

#### 8.4.4 Unsupervised clustering

Unsupervised clustering was performed on the top 50 PCs with Seurat [3] (resolution ∊ {0.8, 1.2}; these represent low-moderate and high levels within the recommended range) and on the `SingleCellExperiment` object of raw and normalized counts with SC3 [2]. We also provided [3] with only the marker genes used by CellAssign (SC3 failed to run when provided with this number of genes). Inferred clusters were mapped to true (FACS-purified) cell types by computing the pairwise Spearman correlation between mean expression vectors for each cluster and each true cell type. Each cluster was treated as the cell type it was most strongly positively associated with by Spearman’s *ρ*.

#### 8.4.5 Evaluation

Accuracy and cell type-level F1 score were computed to evaluate clustering performance. The cell type-level F1 score is defined as the arithmetic mean of F1 scores computed for each cell type separately.

### 8.5 High-grade serous ovarian cancer

#### 8.5.1 Sample preparation

Specimens were placed into cold media in the operating room and brought to the clinical laboratory by messenger porter. Following this, each specimen was assigned a unique research identifier and processed as per VGH/UBC Anatomical Pathology specimen handling procedures. Tissues were dissociated at low temperature [40] using a modified protocol (O’Flanagan et al., manuscript in preparation). Briefly, after finely chopping and weighing in a cell culture dish, tissue was transferred into a gentleMACS C tube, and 1mL of 10 mg/mL *Bacillus lichenformis* protease (Creative Enzymes NATE-0633) was added to each 25 mg of tissue. The resulting solution was incubated and mechanically disrupted at 6^*◦*^C using the Miltenyi Biotec MACS Separator (programs h_tumour_01, h_tumour_02, h_tumour_03) for 1 hour. Following dissociation, cells were assessed for viability using the cell counter (5*μ*L cells + 5*μ*L trypan blue) under a microscope.

Samples were then diluted with cold HFN and washed with trypsin, dispase, and DNAse while gently pipetting up and down. Cold ammonium chloride was added to bloody samples. Cells were assessed for viability using the cell counter (5*μ*L cells + 5*μ*L trypan blue) under a microscope, and kept on ice. Cells were spun down and the pellet resuspended in 100*μ*L of Miltenyi Dead Cell Removal MicroBeads and incubated at room temperature for 15 minutes. Viable cell enrichment was performed using the positive selection column type MS with a MACS Separator.

#### 8.5.2 Library preparation and sequencing

Single-cell RNA-seq libraries were prepared following the 10x Genomics User Guide for 5’ gene expression library construction. Single cell libraries were sequenced on an Illumina NextSeq 500 (75bp paired end reads) using a modified 58bp R2 at the UBC Biomedical Research Centre.

#### 8.5.3 Processing and normalization of single-cell RNA-seq data

Raw sequence files were processed with CellRanger v2.1.0. The resulting filtered count matrices were read into `SingleCellExperiment` objects. Outlier cells according to quality control parameters (≥ 3 median absolute deviations from the median) were filtered out using the `scater` R package. Additionally, cells with ≥ 20% mitochondrial UMIs or ≥ 50% ribosomal UMIs were removed (ovarian cancer cells can have higher mitochondrial percentages than other cell types, as in [41]). Size factors were computed using `quickCluster` and `computeSumFactors` from the `scran` R package. Following this, data normalization was performed using `scater normalize`. Principal components analysis was performed on the resultant normalized logcounts for the top 1000 most variable genes. The first 50 PCs were used as input for UMAP.

For HGSC data, two UMAP parameters were changed from the defaults (`umap` R package) due to the presence of an outlier in UMAP space along the first dimension. The number of neighbours was set to 25, and the minimum distance was set to 0.2.

Cell cycle scores were computed with `cyclone` from the scran package [33, 39].

#### 8.5.4 CellAssign

- B cells: *VIM*^c^, *MS4A1*^c^, *CD79A*^c^, *PTPRC*^c^, *CD19*^c^, *BANK1* [42]
- T cells: *VIM*^c^, *CD2*^c^, *CD3D*^c^, *CD3E* ^c^, *CD3G*^c^, *CD28*^c^, *PTPRC*^c^
- Monocyte/Macrophage: *VIM*^c^, *CD14*^c^, *FCGR3A*^c^, *CD33*^c^, *ITGAX* ^c^, *ITGAM*^c^, *CD4*^c^, *PTPRC*^c^, *LYZ*^c^
- Epithelial cells: *EPCAM*^c^, *CDH1*^c^, *KRT8* [19], *WFDC2* [19]
- Ovarian stromal cells: *VIM*^c^, *MUM1L1* [18], *FOXL2* [18], *ARX* [18], *DCN* [19], *TPT1* [19], *RBP1* [19]
- Ovarian myofibroblast: *VIM*^c^, *MUM1L1* [18], *FOXL2* [18], *ARX* [18], *ACTA2*^c^, *COL1A1*^c^, *COL3A1*^c^, *SERPINH1* [19]
- Vascular smooth muscle cells: *VIM*^c^, *ACTA2*^c^, *MYH11*^c^, *PLN* [43], *MYLK* ^c^, *MCAM* [20], *COL1A1*^c^, *COL3A1*^c^, *SERPINH1* [44]
- Endothelial cells: *VIM*^c^, *EMCN*^c^, *CLEC14A* [45], *CDH5*^c^, *PECAM1*^c^, *VWF* ^c^, *MCAM* [20], *SERPINH1* [19]

^c^: canonical marker

The marker gene list described above and in **Supplemental Table 2** was used for CellAssign [42, 18, 19]. *DCN*, *TPT1*, and *RBP1* were selected as markers of ovarian stromal cells based on differential expression results comparing normal fibroblasts (ovarian stromal cells) and malignant fibroblasts from [19] (these were the top 3 genes upregulated in normal fibroblasts by log fold change where *Q* < 0.05). CellAssign was run with default parameters, the shrinkage prior on *δ*_*gc*_ values turned on, and 5 random initializations.

#### 8.5.5 Unsupervised clustering

Unsupervised clustering of epithelial cells from CellAssign (probability ≥ 90%) was performed with Seurat [3], using a resolution parameter of 0.2 (for fairly coarse resolution). Unsupervised clustering of all cells was performed with Seurat and SC3 [2], using default parameters. For Seurat, resolutions of 0.8 and 1.2 were used (these represent low-moderate and high levels within the recommended range). Additionally, Seurat clustering was also performed using data for the same set of marker genes provided to CellAssign (SC3 failed to run when provided with this number of genes).

#### 8.5.6 Differential expression

Log fold change values from the `findMarkers` function (filtering out ribosomal and mitochondrial genes) from `scran` were used as input for gene set enrichment analysis with the fGSEA R package, using default parameters with 10,000 permutations, and the hallmark pathway gene set [32]. Annotations for cell cycle-associated pathways (E2F targets, G2M checkpoint, and mitotic spindle), and immune-associated pathways (including interferon gamma response, interferon alpha response, coagulation, complement, IL6-JAK/STAT signalling, and allograft rejection) were taken from [32].

### 8.6 Follicular lymphoma

#### 8.6.1 Sample preparation

Leftovers from clinical flowed samples were collected and frozen in fetal calf serum containing 10% DMSO. Cells were thawed and washed according to the steps outlined in the 10X Genomics Sample Preparation Protocol. Cells were stained with PI for viability and sorted in a BD FACSAria Fusion using a 85um nozzle. Sorted cells were collected in 0.5 ml of medium, centrifuged and diluted in 1X PBS with 0.04% bovine serum albumin.

#### 8.6.2 Library preparation and sequencing

Cell concentration was determined by using a Countess II Automated Cell Counter and approximately 3,500 cells were loaded per well in the Single Cell 3’ Chip. Single cell libraries were prepared according to the Chromium Single Cell 3’Reagent Kits V2 User Guide. Single cell libraries from two samples were pooled and sequenced on one HiSeq 2500 125 base PET lane.

#### 8.6.3 Preprocessing and normalization of single-cell RNA-seq data

Preprocessing steps for the follicular lymphoma data were identical to those for HGSC single-cell RNA-seq data, described in section 8.5.3, with the exception of different mitochondrial and ribosomal thresholds (cells with ≥ 10% mitochondrial UMIs or ≥ 60% ribosomal UMIs were removed).

#### 8.6.4 scvis analysis

scvis train (v0.1.0) [26] was run with default settings on the top 50 PCs to produce a 2-dimensional embedding of the follicular lymphoma data. Early stopping was added to scvis, so that the model would terminate after 3 successive iterations of no improvement (relative improvement in ELBO < 10^−5^). The resultant model was saved and used for mapping in section 8.7.4.

#### 8.6.5 CellAssign

- B cells: *CD19*^c^, *MS4A1*^c^, *CD79A*^c^, *CD79B*^c^, *CD74*^c^, *CXCR5* [46]
- Cytotoxic T cells: *CD2*^c^, *CD3D*^c^, *CD3E* ^c^, *CD3G*^c^, *TRAC*^c^, *CD8A*^c^, *CD8B*^c^, *GZMA*^c^, *NKG7* ^c^, *CCL5*^c^, *EOMES*^c^
- Follicular helper T cells: *CD2*^c^, *CD3D*^c^, *CD3E* ^c^, *CD3G*^c^, *TRAC*^c^, *CD4*^c^, *CXCR5*^c^, *PDCD1*^c^, *TNFRSF4* [42], *ST8SIA1* [42], *ICA1* [42], *ICOS* [42]
- Other CD4+ T cells: *CD2*^c^, *CD3D*^c^, *CD3E* ^c^, *CD3G*^c^, *TRAC*^c^, *CD4*^c^, *IL7R* [42]

^c^: canonical marker

The marker gene list described above and in **Supplemental Table 2** was used for CellAssign [42, 47]. CellAssign was run with default parameters, the shrinkage prior on *δ*_*gc*_ values turned on, and 5 random initializations.

Patient was added as an additional covariate into the design matrix *X* (section 8.2.1). The best result according to marginal log-likelihood at convergence was kept. Optimization was considered converged after 3 consecutive rounds of no improvement (relative change in log-likelihood < 10^−5^). MAP assignments from CellAssign were used for downstream analysis.

No evidence of regulatory T cells (*FOXP3* and *IL2RA* expression), NK cells (*NCAM1* expression), and myeloid cells (*CD14*/*CD16* and *LYZ* expression) was detected.

#### 8.6.6 Unsupervised clustering

Unsupervised clustering of all cells was performed with Seurat and SC3 [2], using default parameters. For Seurat, resolutions of 0.8 and 1.2 were used (these represent low-moderate and high levels within the recommended range). Additionally, Seurat clustering was also performed using data for the same set of marker genes provided to CellAssign (SC3 failed to run when provided with this number of genes).

#### 8.6.7 Classifying B cells

B cells from CellAssign were further subclassified into ‘malignant’ or ‘nonmalignant’ groups according to expression of the constant region of the immunoglobulin light chain (kappa or lambda type) and the results of PCA. Seurat [3] (resolution = 0.8) was used to separate B cells into clusters, based on the top 50 PCs. Following this, the sole cluster associated with *IGKC* (immunoglobulin light chain kappa-type constant region) expression was designated as nonmalignant. We further reasoned this was the case based on the cluster containing a mixture of T1 and T2 cells and constituting only a minor subset of the B cells.

#### 8.6.8 Differential expression between timepoints

Differential expression analysis between timepoints for a given celltype and patient was performed using voom from the limma package for each patient and cell type separately, with timepoint as the independent variable. Genes with low expression (< 500 UMIs in total across all cells) were removed. *P*-values were adjusted with the Benjamini-Hochberg method, and genes with *Q* ≤ 0.05 were considered differentially expressed. Differential expression between malignant and nonmalignant B cells was performed similarly, but using the formula `~malignant_status + timepoint + malignant_status:timepoint` to control for timepoint and any interactions.

#### 8.6.9 Reactome pathway enrichment analysis

Pathway analysis was performed for the top 50 most upregulated and top 50 most downregulated genes (separately) by log fold change from limma (where *Q* ≤ 0.05, filtering out ribosomal and mitochondrial genes). Overrepresentation of Reactome [30] pathways was assessed using the R package ReactomePA. Pathways were considered significantly overrepresented if the adjusted *P*-value ≤ 0.05 and at least 2 genes from the pathway were present.

#### 8.6.10 Comparing malignant cells between timepoints

Log fold change values from the `findMarkers` function (filtering out ribosomal and mitochondrial genes) from `scran` were used as input for gene set enrichment analysis with the fGSEA R package, using default parameters with 10,000 permutations, and the hallmark pathway gene set [32]. Annotations for cell cycle-associated pathways (E2F targets, G2M checkpoint, and mitotic spindle) and proliferation-associated pathways were taken from [32]. BH-adjusted *P*-values for differences in replication-associated marker expression (*MKI67* and *TOP2A*) were also computed with the `findMarkers` function from `scran`, using default parameters.

### 8.7 Reactive lymph node data

#### 8.7.1 Sample preparation

Cell suspensions from patients with reactive lymphoid hyperplasia but no evidence of malignant disease and collagen disease were used. Leftovers from clinical flowed samples were collected and frozen in FCS containing 10%DMSO. The day of the experiment cell suspensions were rapidly thawed at 37°C, and washed according to the steps outlined in the 10X Genomics Sample Preparation Protocol. Cells were stained with DAPI and viable cells (DAPI negative) were sorted on a FACS ARIAIII or FACS Fusion (BD Biosciences) instrument.

#### 8.7.2 Library preparation and sequencing

Approximately 8,700 cells per sample were loaded into a Chromium Single Cell 3’ Chip kit v2 (PN-120236) and processed according to the Chromium Single Cell 3’Reagent kit v2 User Guide. Libraries were constructed using the Single 3’ Library and Gel Bead Kit v2 (PN-120237) and Chromium i7 Mulitiplex Kit v2 (PN-120236). Single cell libraries from two samples were pooled and sequenced on one HiSeq 2500 125 base PET lane.

#### 8.7.3 Preprocessing and normalization of single cell RNA-seq data

Preprocessing steps for the reactive lymph node data were identical to those for follicular single-cell RNA-seq data, described in section 8.6.3.

#### 8.7.4 scvis analysis

The identities of the top 1000 most variable genes and PCA loadings from follicular lymphoma data analysis were used to compute a 50-dimensional embedding for the reactive lymph node data. Following this, the resultant 50 PCs were provided as input to scvis map [26], using the model trained in section 8.6.4 and default settings.

## List of Supplementary Figures

**Supplemental Figure 1.**
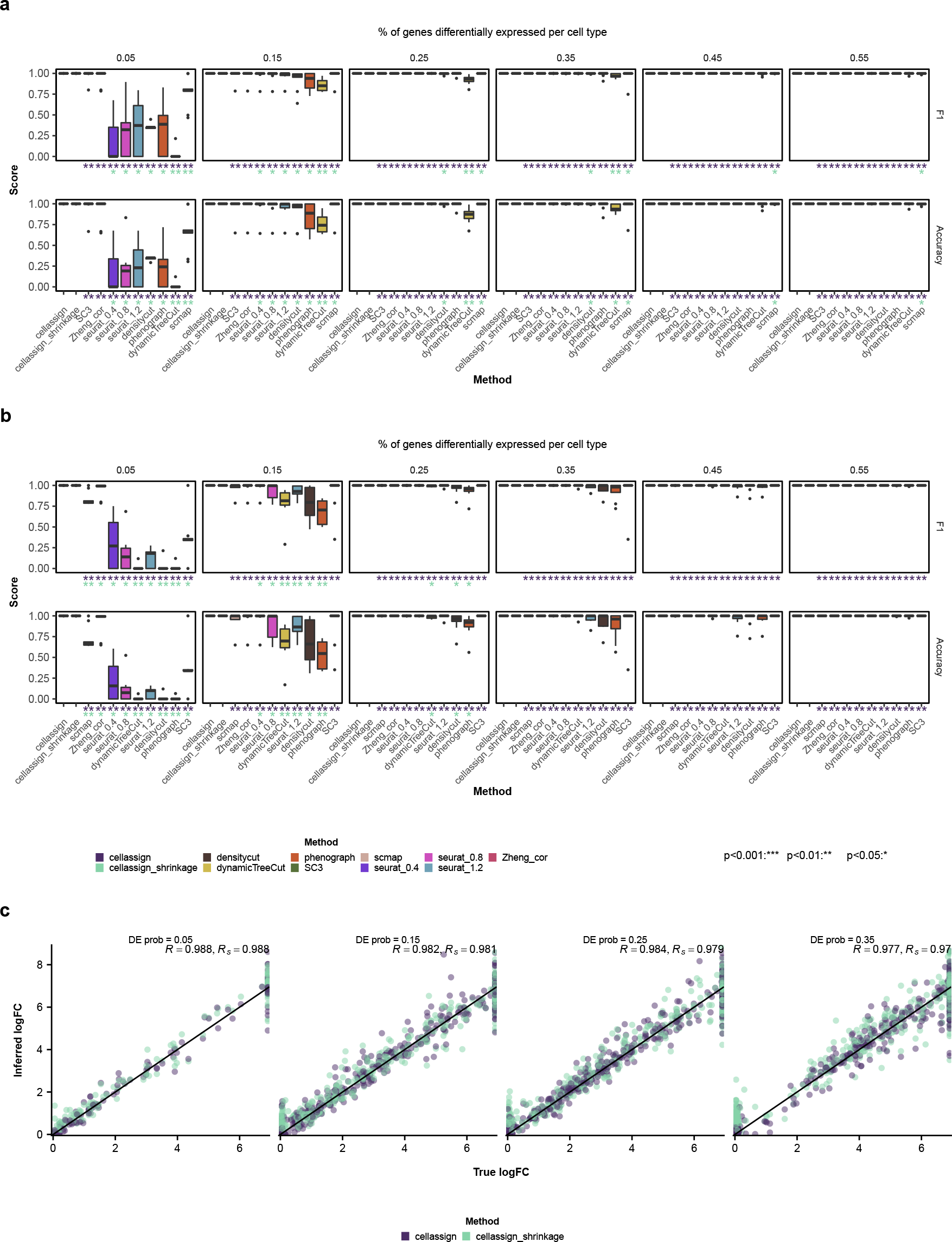
Simulation performance across a range of proportions of differentially expressed genes, using differential expression parameters derived from comparing B and CD8+ T cells. (**a**) Accuracy and cell-level F1 score (**Methods**) for varying proportions of differentially expressed genes per cell type. CellAssign was provided with a set of marker genes (**Methods**); all other methods were provided with all genes. Asterisks indicate FDR-adjusted statistical significance (Wilcoxon signed-rank test) for pairwise comparisons between cellassign/cellassign_shrinkage and other methods. (**b**) Accuracy and cell-level F1 score for varying proportions of differentially expressed genes per cell type. All methods were provided with the same set of marker genes. (**c**) Correspondence between true simulated log fold change values and log fold change (*δ*) values inferred by CellAssign. *R* and *R*_*s*_ refer to the Pearson correlation between true and inferred logFC values for cellassign and cellassign_shrinkage, respectively.

**Supplemental Figure 2.**
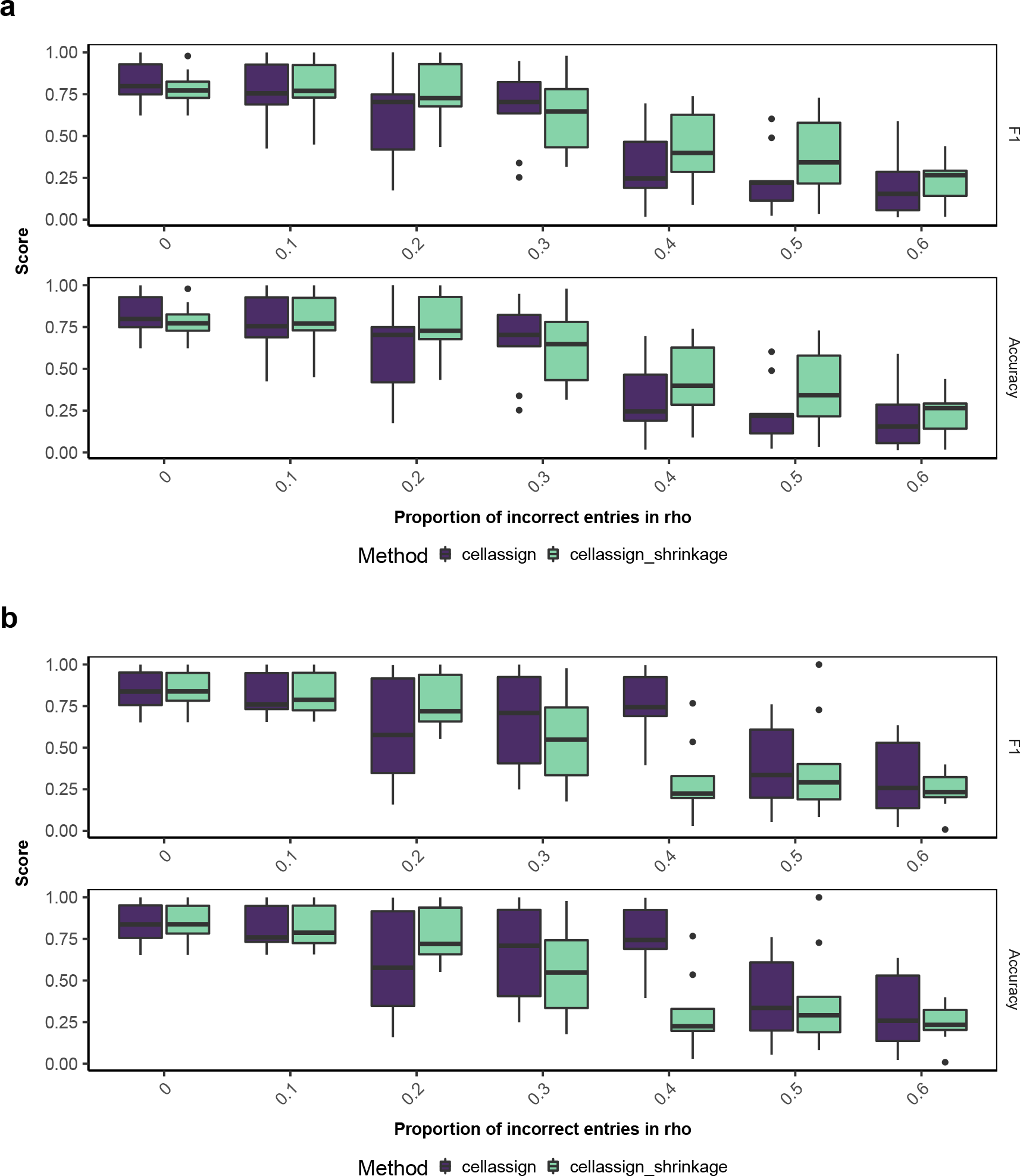
Simulation performance across a range of proportions of randomly flipped entries in the binary marker gene matrix, using differential expression parameters derived from comparing naïve CD8+ and naïve CD4+ T cells for (**a**) 5 marker genes per cell type and (**b**) 20 marker genes per cell type.

**Supplemental Figure 3.**
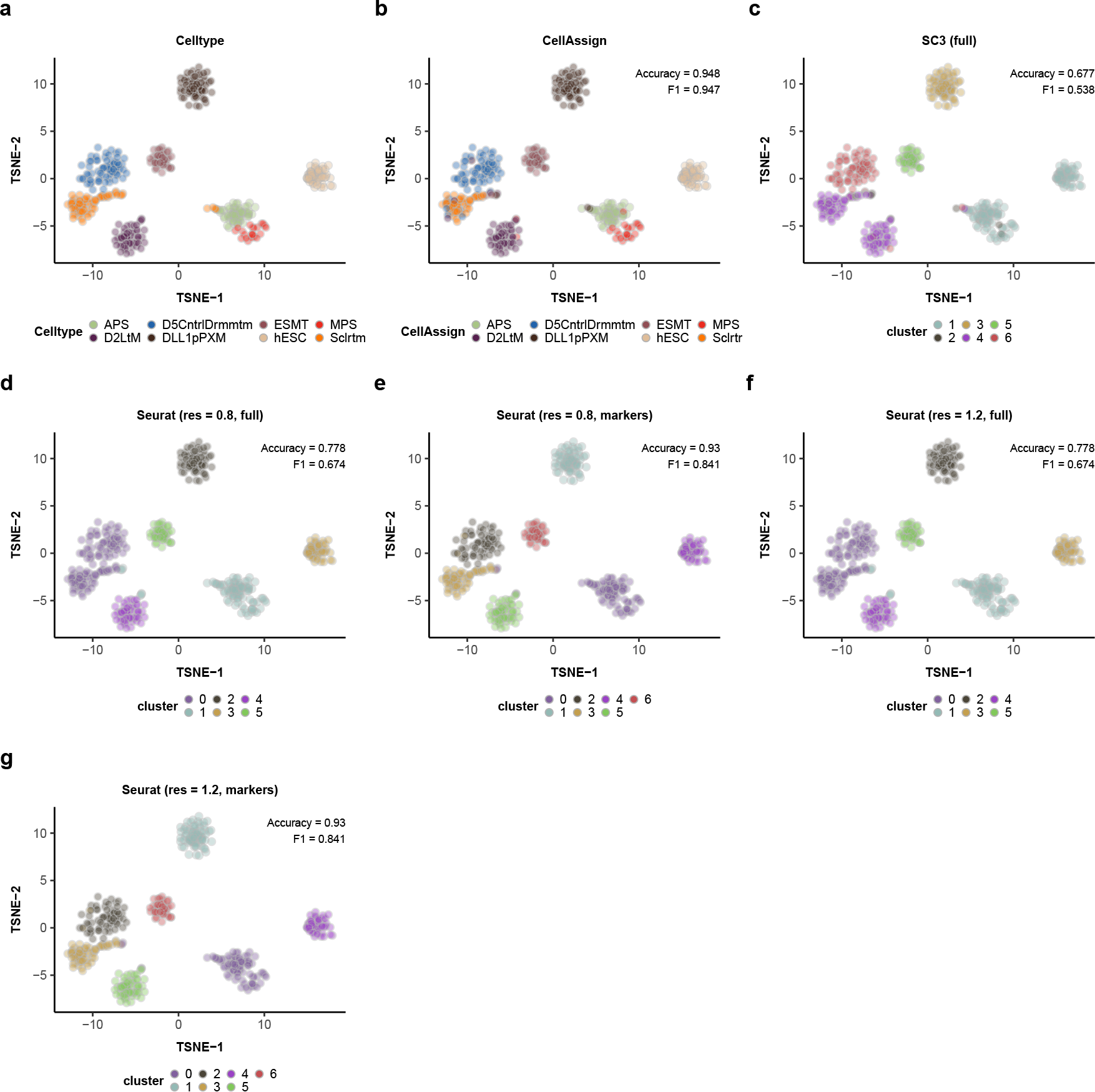
Performance (accuracy and cell type-level F1 score, **Methods**) of CellAssign and the best-performing clustering methods evaluated by [6] on FACS-purified H7 human embryonic stem cells in various stages of differentiation. t-SNE plots of (**a**) ground-truth FACS annotations; (**b**) CellAssign-derived annotations; (**c**) SC3 clusters (using all genes); (**d**) Seurat clusters (resolution = 0.8, using all genes); (**e**) Seurat clusters (resolution = 0.8, using the same marker gene set used by CellAssign); (**f**) Seurat clusters (resolution = 1.2, using all genes); (**g**) Seurat clusters (resolution = 1.2, using the same marker gene set used by CellAssign).

**Supplemental Figure 4.**
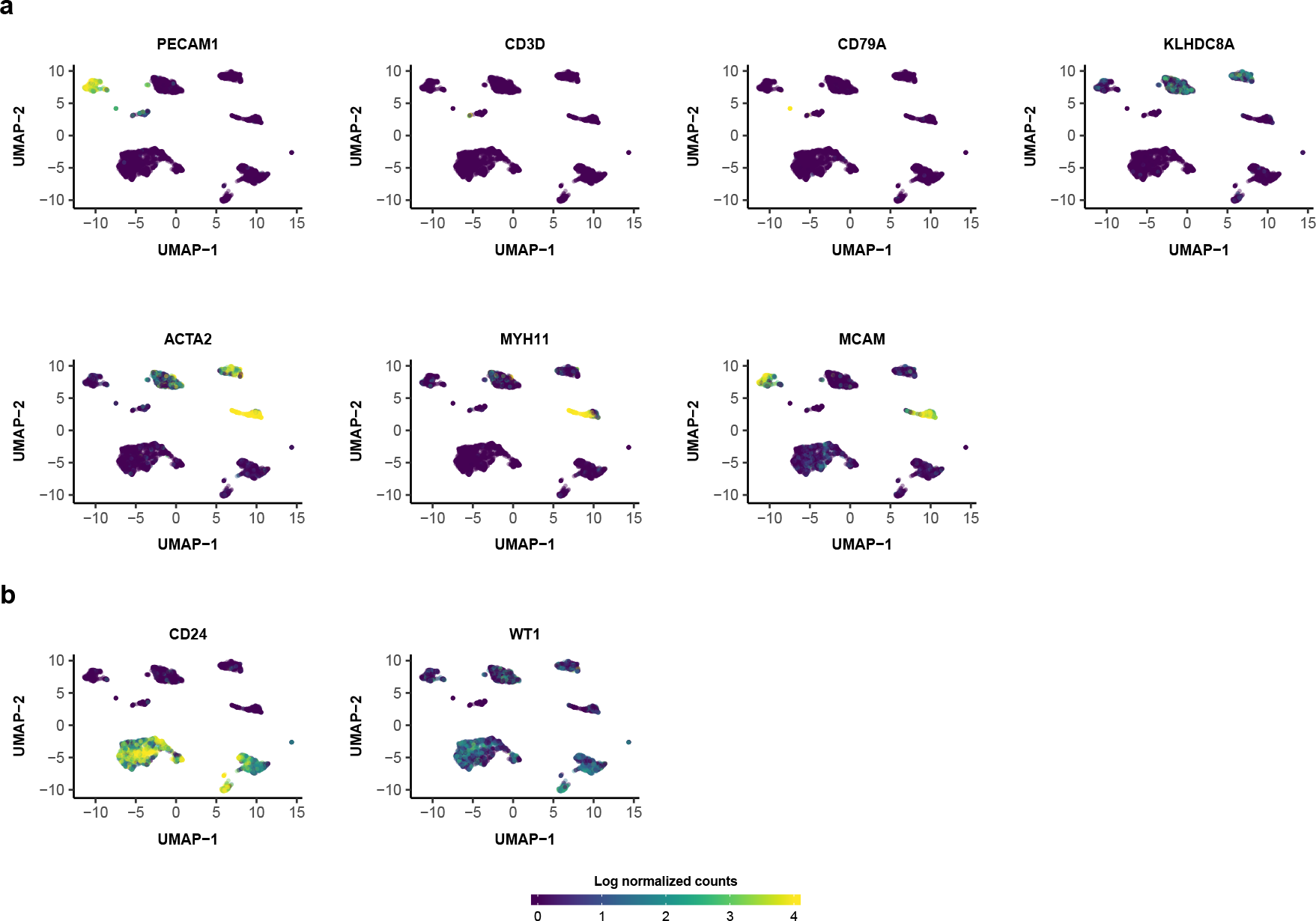
Expression of select marker genes in HGSC single cell RNA-seq data. (**a**) Expression (log normalized counts) of *PECAM1* (for endothelial cells), *CD3D* (for T cells), *CD79A* (for B cells), *KLHDC8A* (for ovary-derived cells), *ACTA2* (for myofibroblasts and smooth muscle), *MYH11* (for smooth muscle), and *MCAM* (for vascular cell types including endothelial cells, vascular smooth muscle, and pericytes). (**b**) Expression (log normalized counts) of marker genes expressed in epithelial ovarian cancers but not in normal ovarian tissue. Expression values were winsorized between 0 and 4.

**Supplemental Figure 5.**
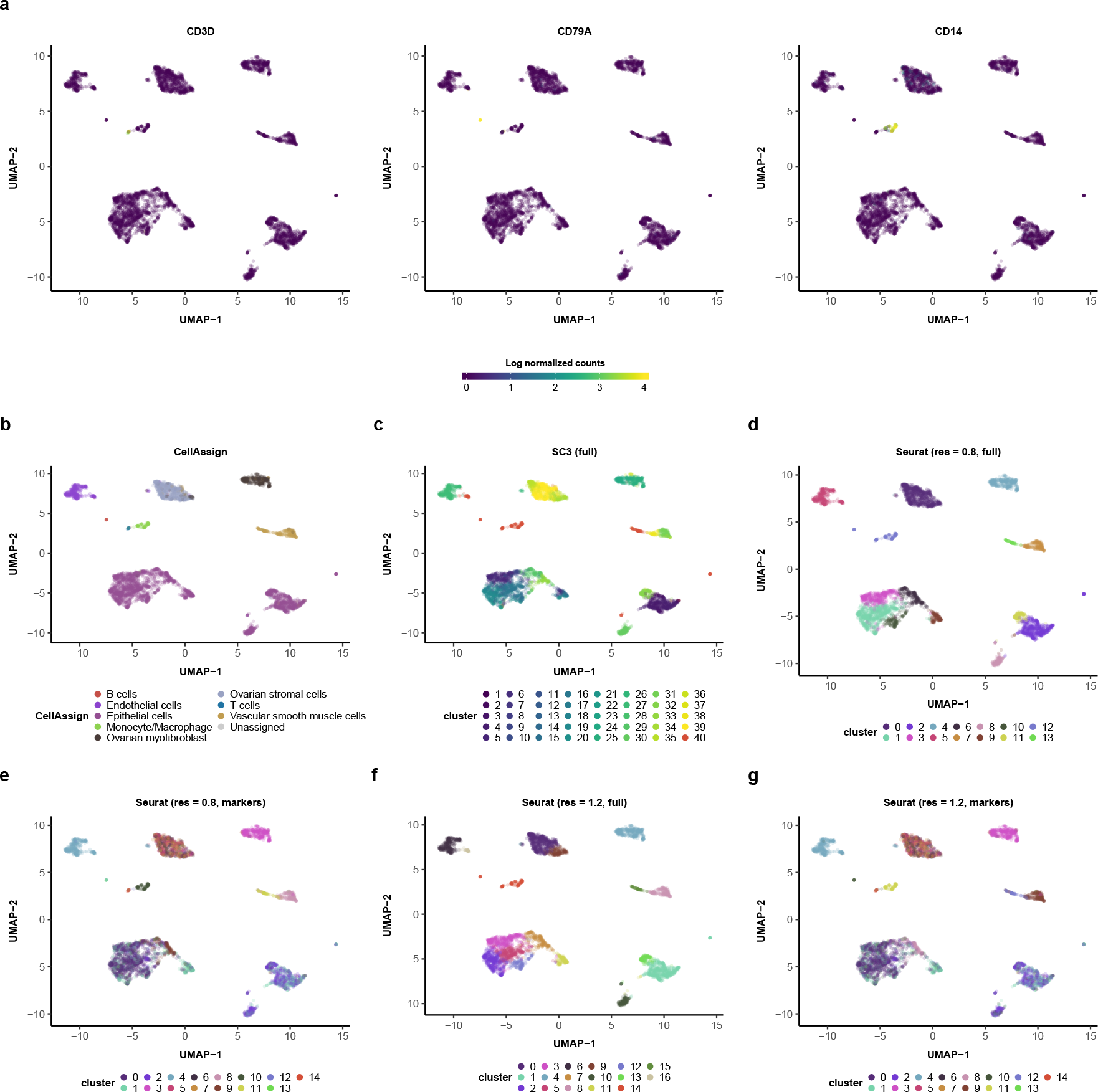
Comparison of clusters from CellAssign and state-of-the-art unsupervised clustering approaches [6] on HGSC single cell RNA-seq data. (**a**) Expression (log normalized counts) of key marker genes of hematopoietic subpopulations *CD3D* (for T cells), *CD79A* (for B cells), and *CD14* (for monocytes/macrophages). Expression values were winsorized between 0 and 4. UMAP plots of (**b**) CellAssign-derived annotations; (**c**) SC3 clusters (using all genes); (**d**) Seurat clusters (resolution = 0.8, using all genes); (**e**) Seurat clusters (resolution = 0.8, using the same marker gene set used by CellAssign); (**f**) Seurat clusters (resolution = 1.2, using all genes); (**g**) Seurat clusters (resolution = 1.2, using the same marker gene set used by CellAssign).

**Supplemental Figure 6.**
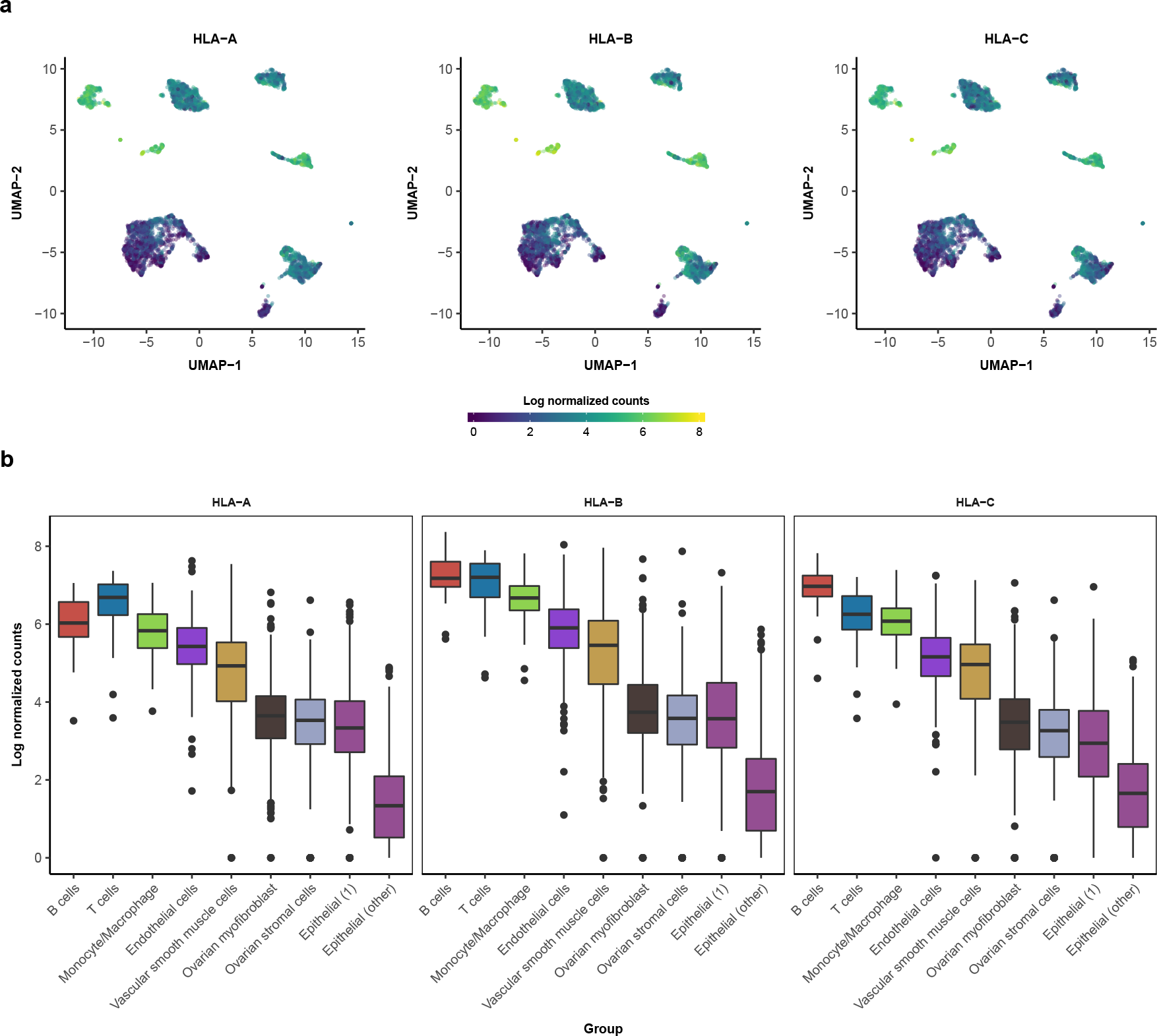
Cluster-specific HLA expression in HGSC epithelial cells. (**a**) Expression (log normalized counts) of HLA class I genes in all HGSC cells. Expression values clipped from 0 to 8. (**b**) Expression of HLA class I genes across cell types in all HGSC cells. Epithelial (1): epithelial cells from cluster 1. Epithelial (other): epithelial cells from all other clusters.

**Supplemental Figure 7.**
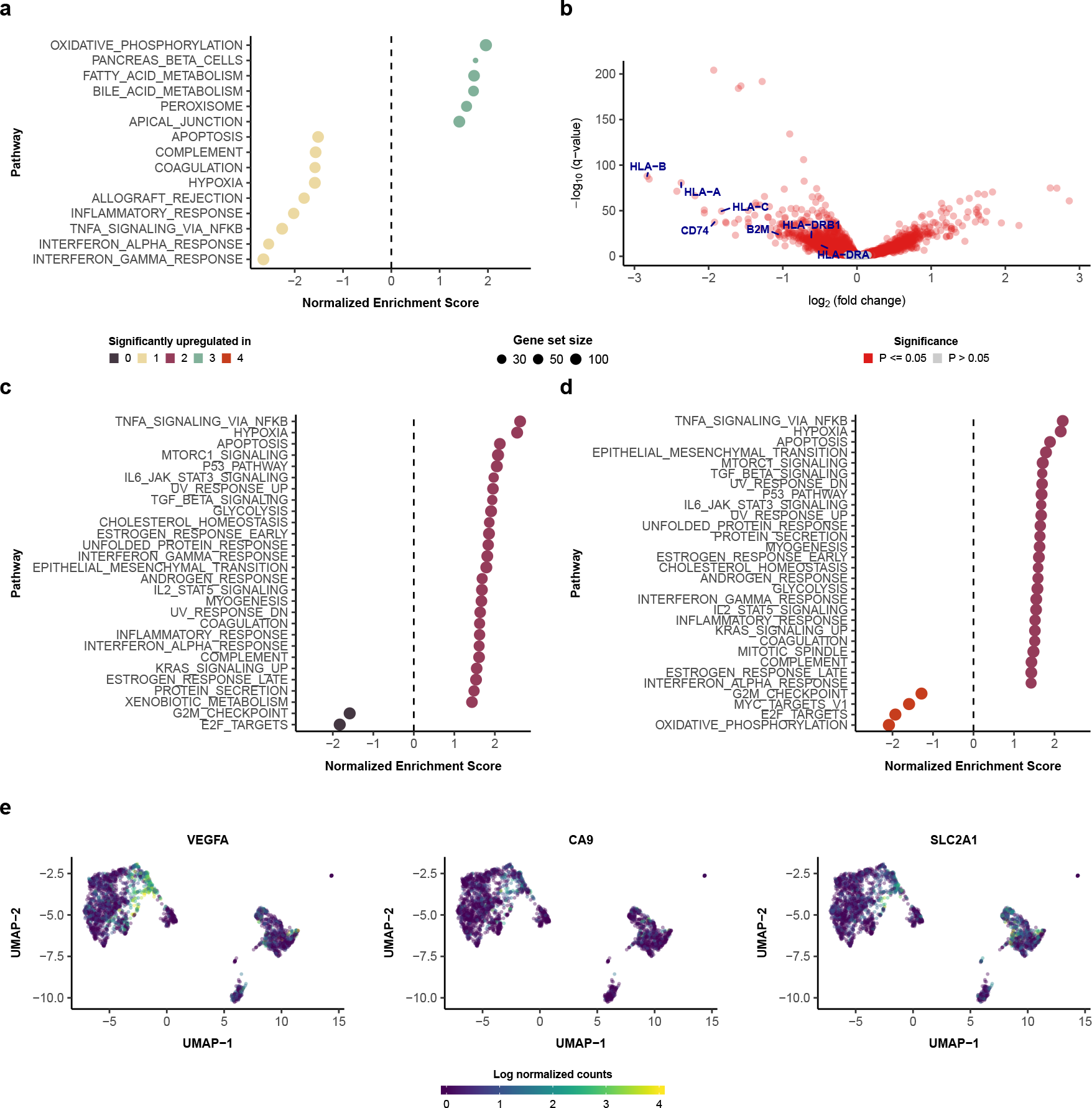
Cluster-specific phenotypes in HGSC epithelial cells. (**a**) Hallmark pathway enrichment results for epithelial clusters 3 vs. 1 from the left ovary sample. (**b**) Gene-level differential expression for epithelial clusters 3 vs. 1. (**c**and **d**) Hallmark pathway enrichment results for epithelial clusters (**c**) 2 vs. 0; and (**d**) 2 vs. 4. (**e**) Expression (log normalized counts) of select hypoxia-associated markers in HGSC epithelial cells. Expression values were winsorized between 0 and 4.

**Supplemental Figure 8.**
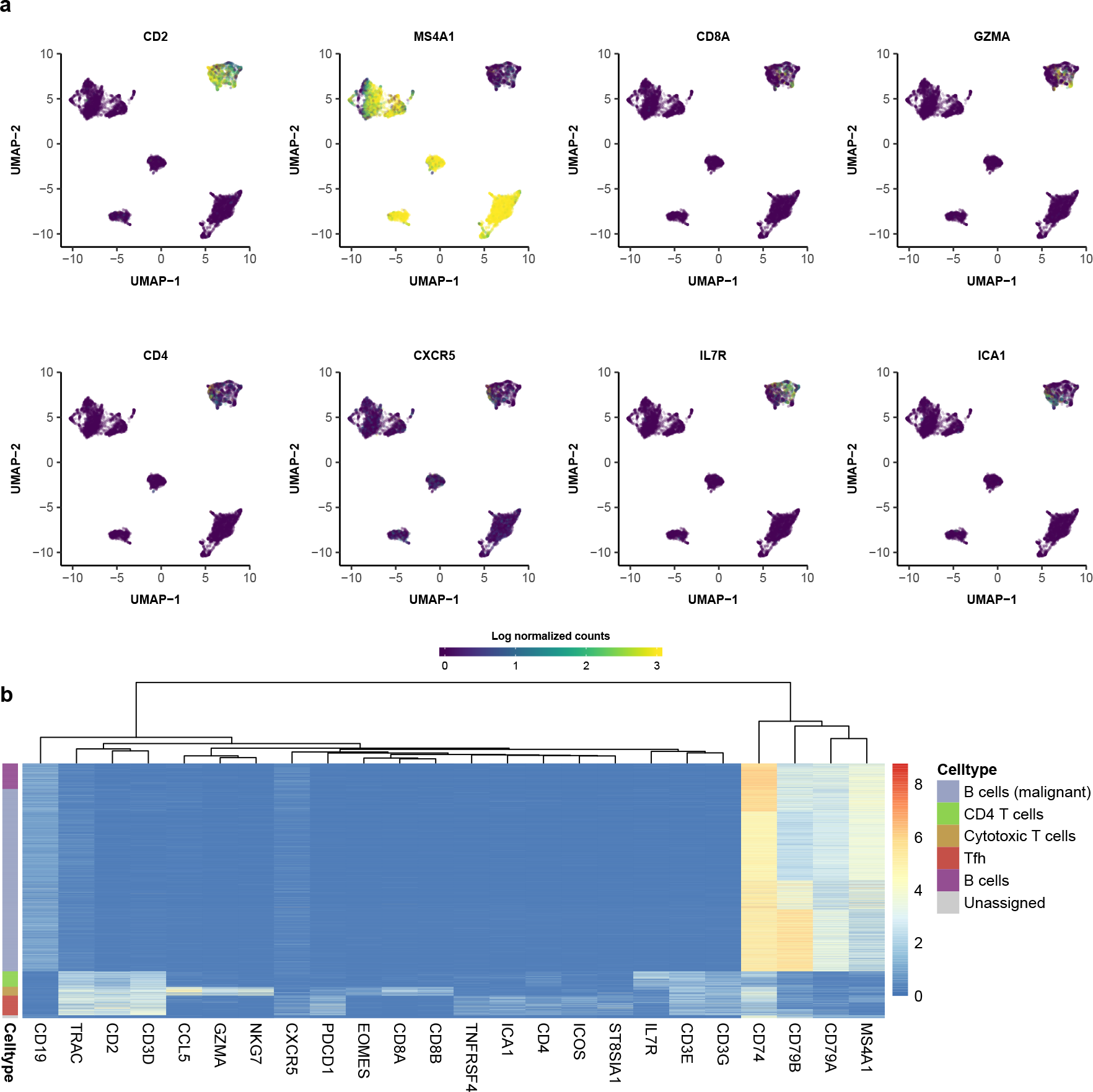
(**a**) Expression (log normalized counts) of select marker genes *CD2* (for T cells), *MS4A1* (for B cells), *CD8A* and *GZMA* (for CD8+ T cells), *CD4* (for T follicular helper cells and other CD4+ T cells) and *CXCR5* and *ICOS* (for T follicular helper cells). Expression values were winsorized between 0 and 3. (**b**) Heatmap of marker gene expression, labeled by maximum probability CellAssign-inferred cell types.

**Supplemental Figure 9.**
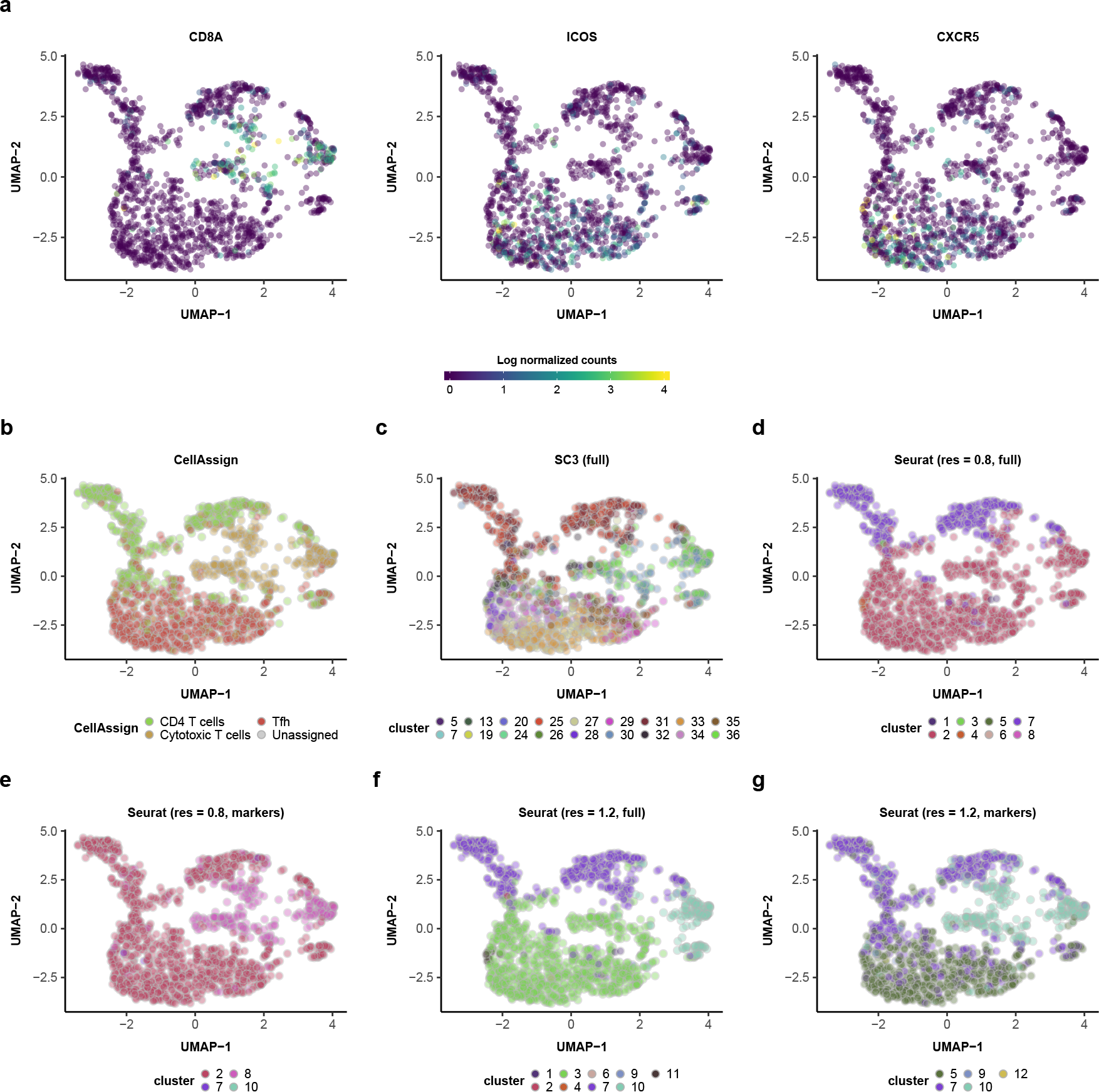
Comparison of clusters from CellAssign and state-of-the-art unsupervised clustering approaches [6] on follicular lymphoma single cell RNA-seq data (showing only T cell subtypes). (**a**) Expression (log normalized counts) of key T cell subpopulation marker genes *CD8A* (for cytotoxic T cells), *ICOS* and *CXCR5* (for T follicular helper cells). Expression values were winsorized between 0 and 4. UMAP plots of (**b**) CellAssign-derived annotations; (**c**) SC3 clusters (using all genes); (**d**) Seurat clusters (resolution = 0.8, using all genes); (**e**) Seurat clusters (resolution = 0.8, using the same marker gene set used by CellAssign); (**f**) Seurat clusters (resolution = 1.2, using all genes); (**g**) Seurat clusters (resolution = 1.2, using the same marker gene set used by CellAssign).

**Supplemental Figure 10.**
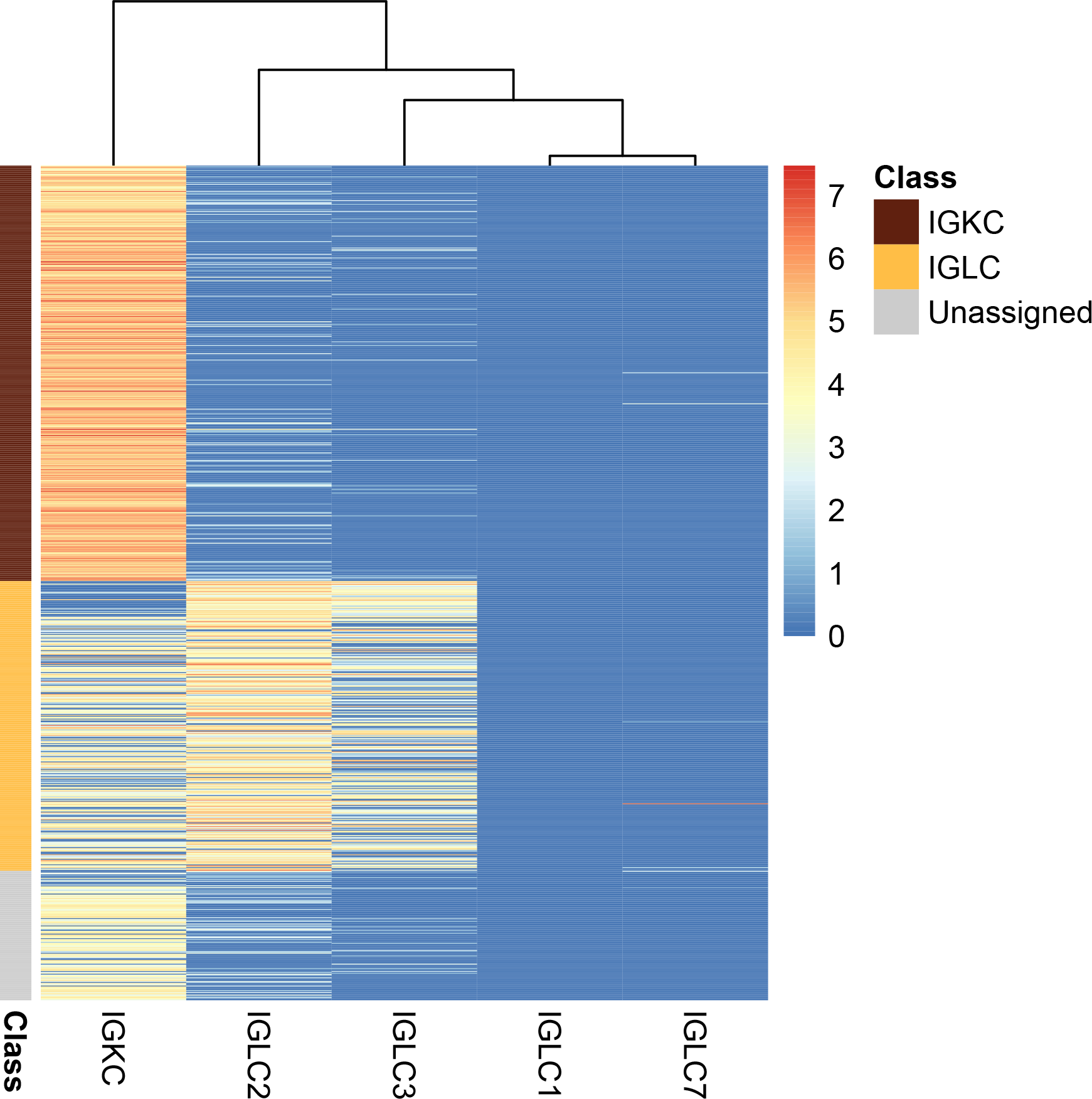
Expression (log normalized counts) of *κ* and *λ* light chain constant region genes in nonmalignant B cells. Class assignments were determined by CellAssign (**Methods**).

**Supplemental Figure 11.**
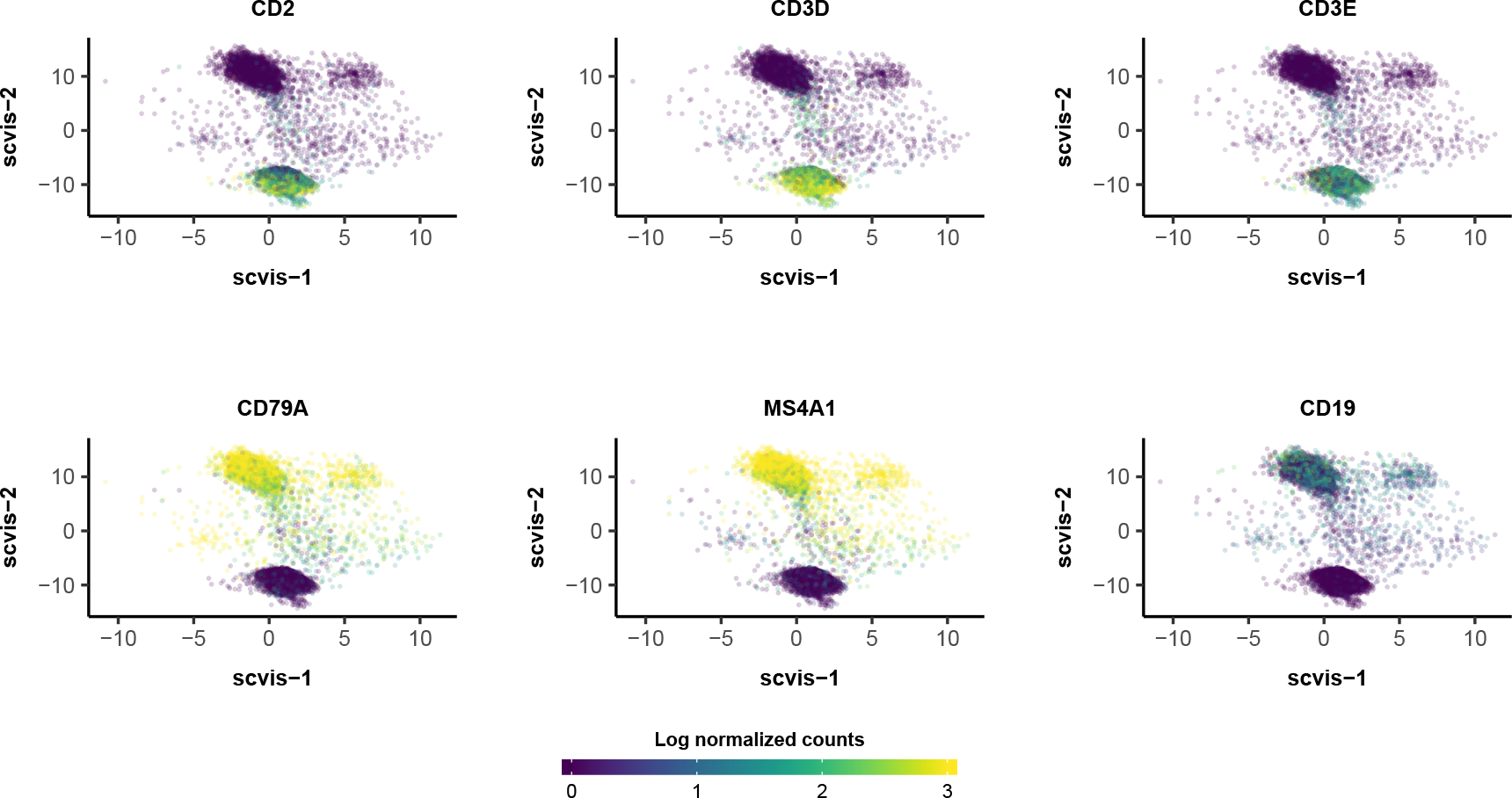
Expression (log normalized counts) of selected marker genes (*CD2*, *CD3D*, and *CD3E* for T cells; *CD79A*, *MS4A1*, and *CD19* for B cells) in scvis embedding of reactive lymph node data. Expression values were winsorized between 0 and 3.

**Supplemental Figure 12.**
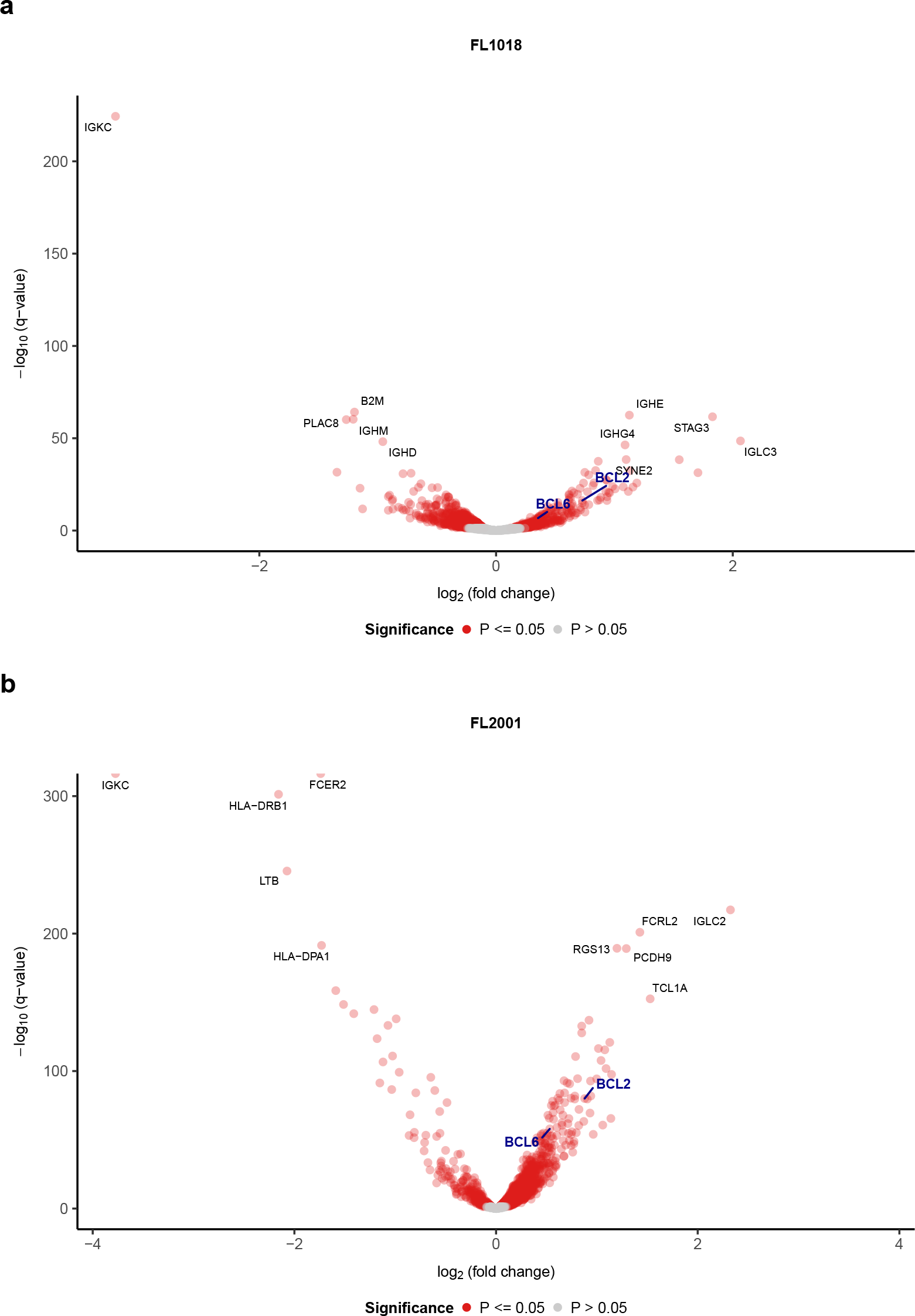
Differential expression results for malignant vs. nonmalignant B cells in (**a**) FL1018 and (**b**) FL2001. Comparisons was performed accounting for timepoint and potential interactions between malignant status and timepoint using a multivariate linear model described in **Methods**. Genes upregulated among malignant cells have logFC values > 0. *P*-values were adjusted with the Benjamini-Hochberg method.

**Supplemental Figure 13.**
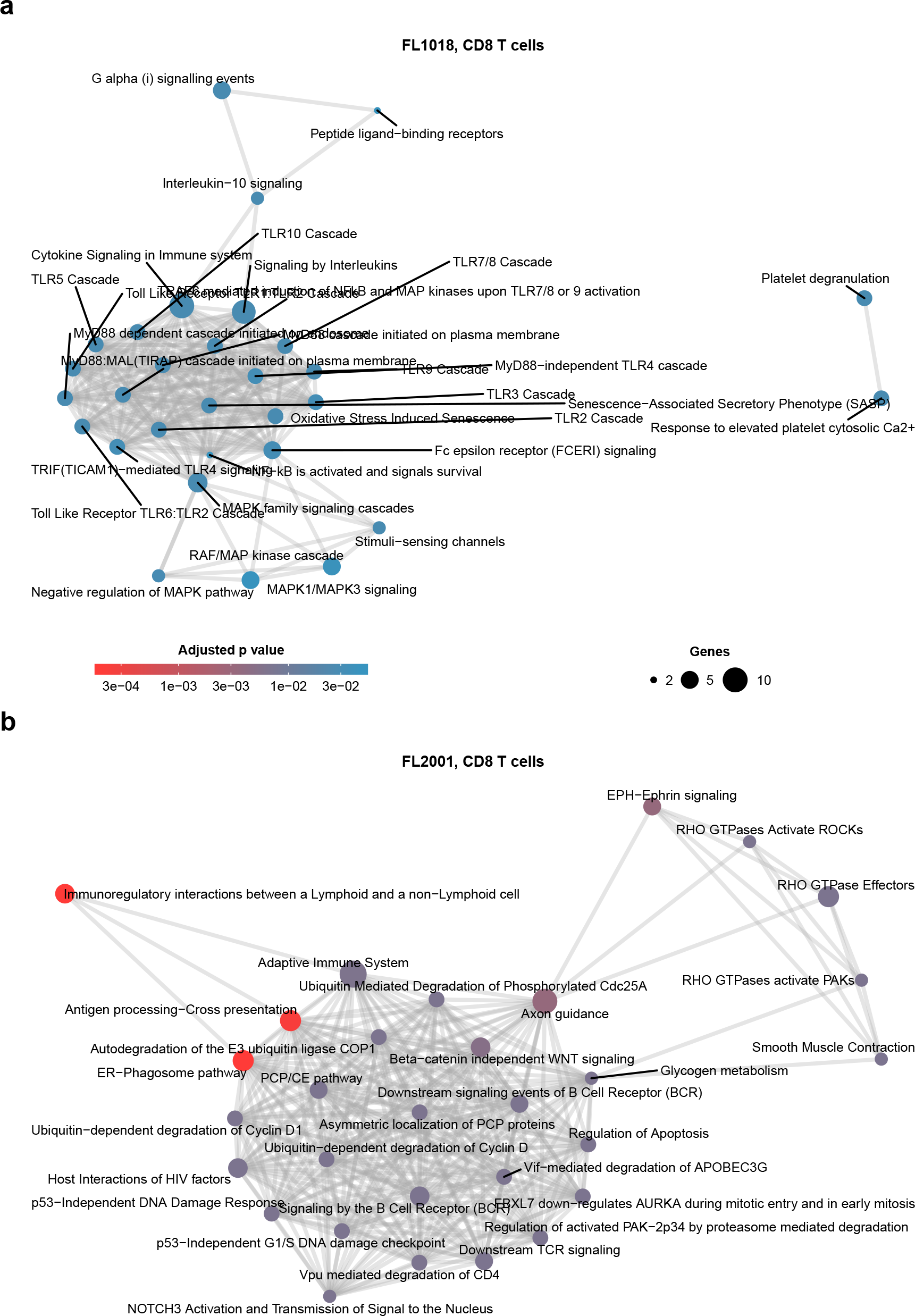
Significantly enriched Reactome pathways (BH-adjusted *P* -value ≤ 0.05) among the top 50 most highly upregulated genes (ranked by log fold change) in (**a**) FL1018 and (**b**) FL2001. Up to 30 pathways are shown in either plot (**Methods**).

**Supplemental Figure 14.**
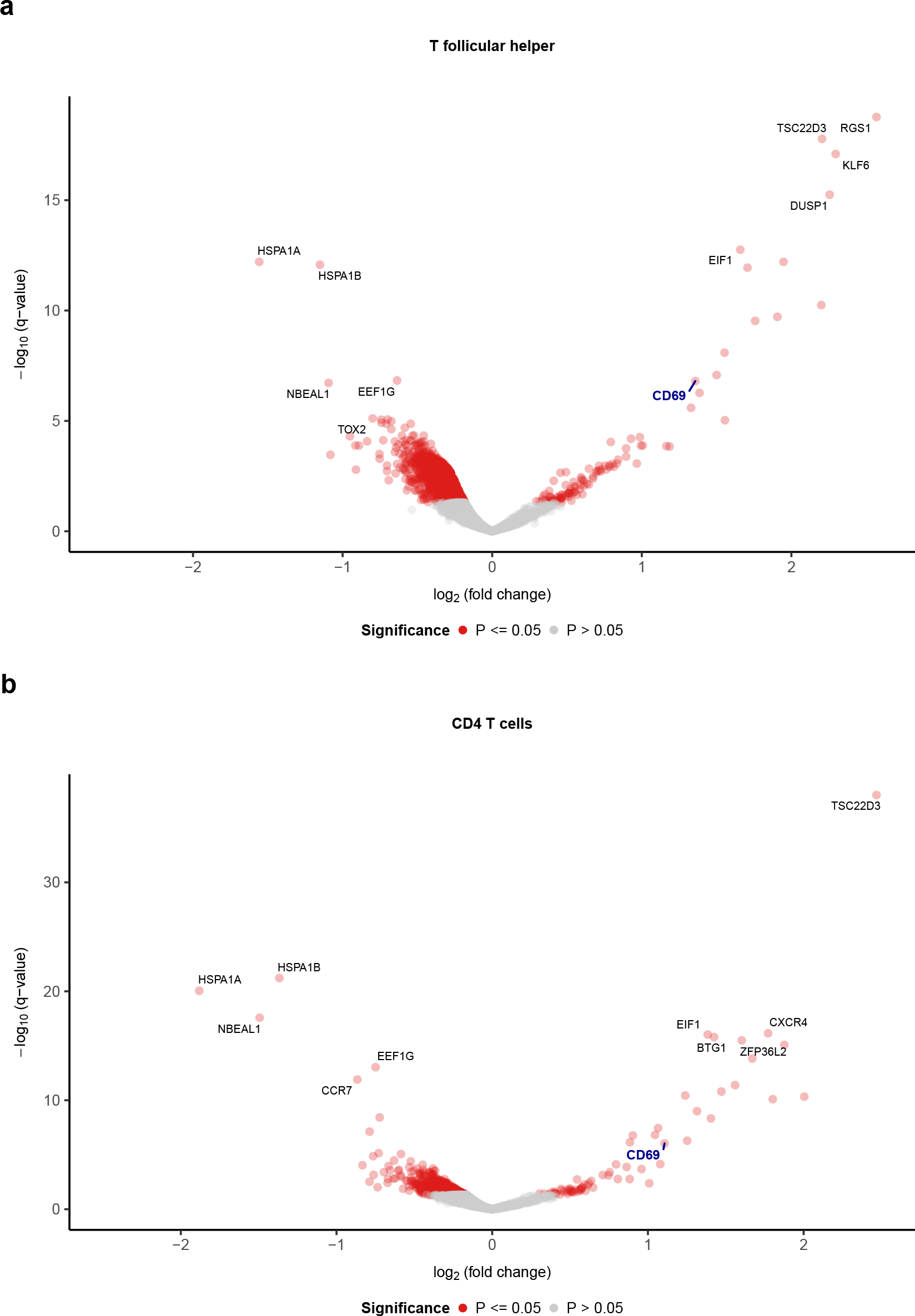
Differentially expressed genes for (**a**) T follicular helper and (**b**) other CD4 T cells between T2 vs. T1. Genes upregulated in T2 have log fold change values > 0. The activation marker *CD69* is highlighted. *P*-values were adjusted with the Benjamini-Hochberg method.

**Supplemental Figure 15.**
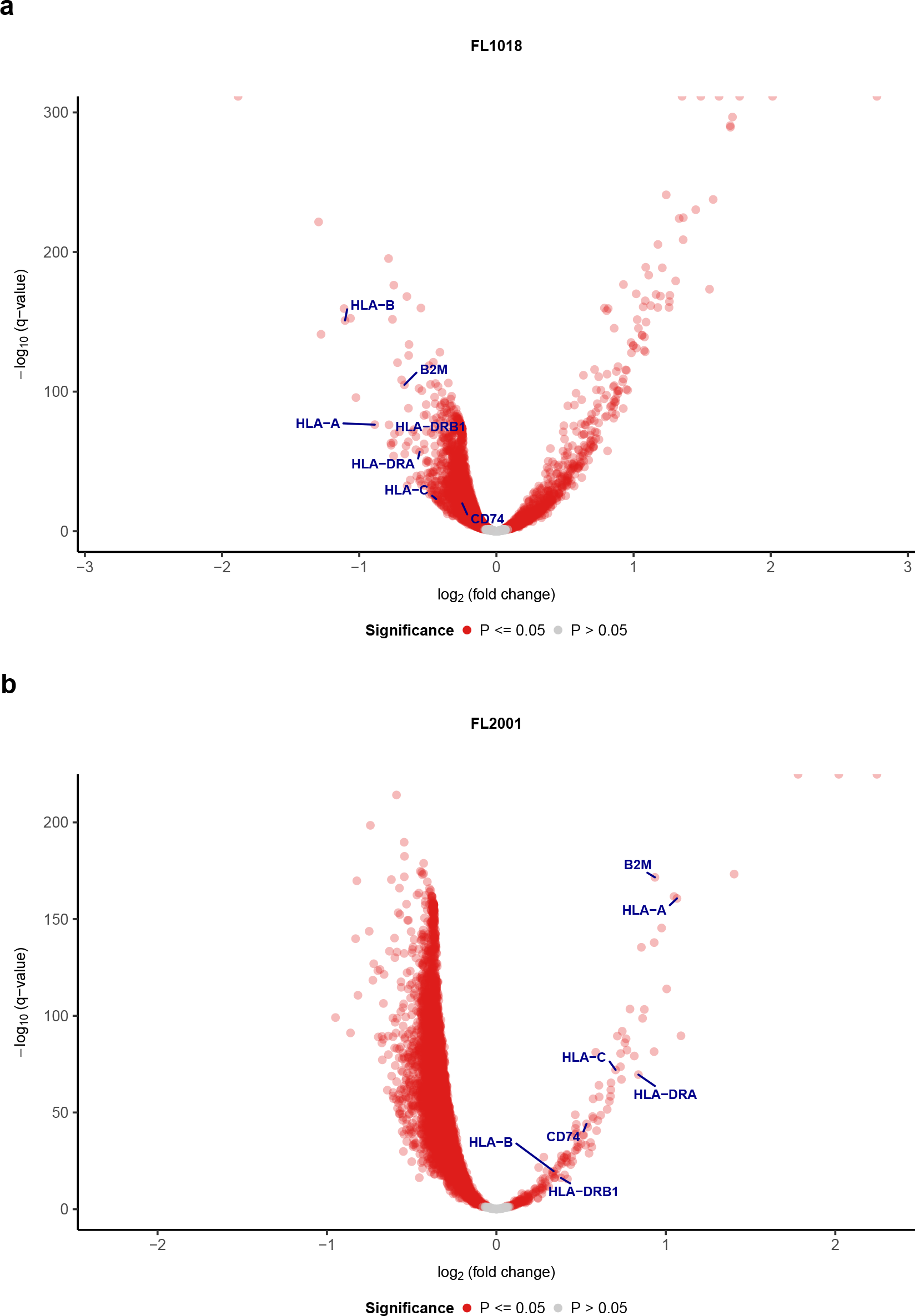
Differentially expressed genes between malignant B cells from T2 vs. T1 in (**a**) FL1018 and (**b**) FL2001. Genes upregulated in T2 have log fold change values > 0. HLA class I genes and select HLA class II genes are highlighted. *P*-values were adjusted with the Benjamini-Hochberg method.

**Supplemental Figure 16.**
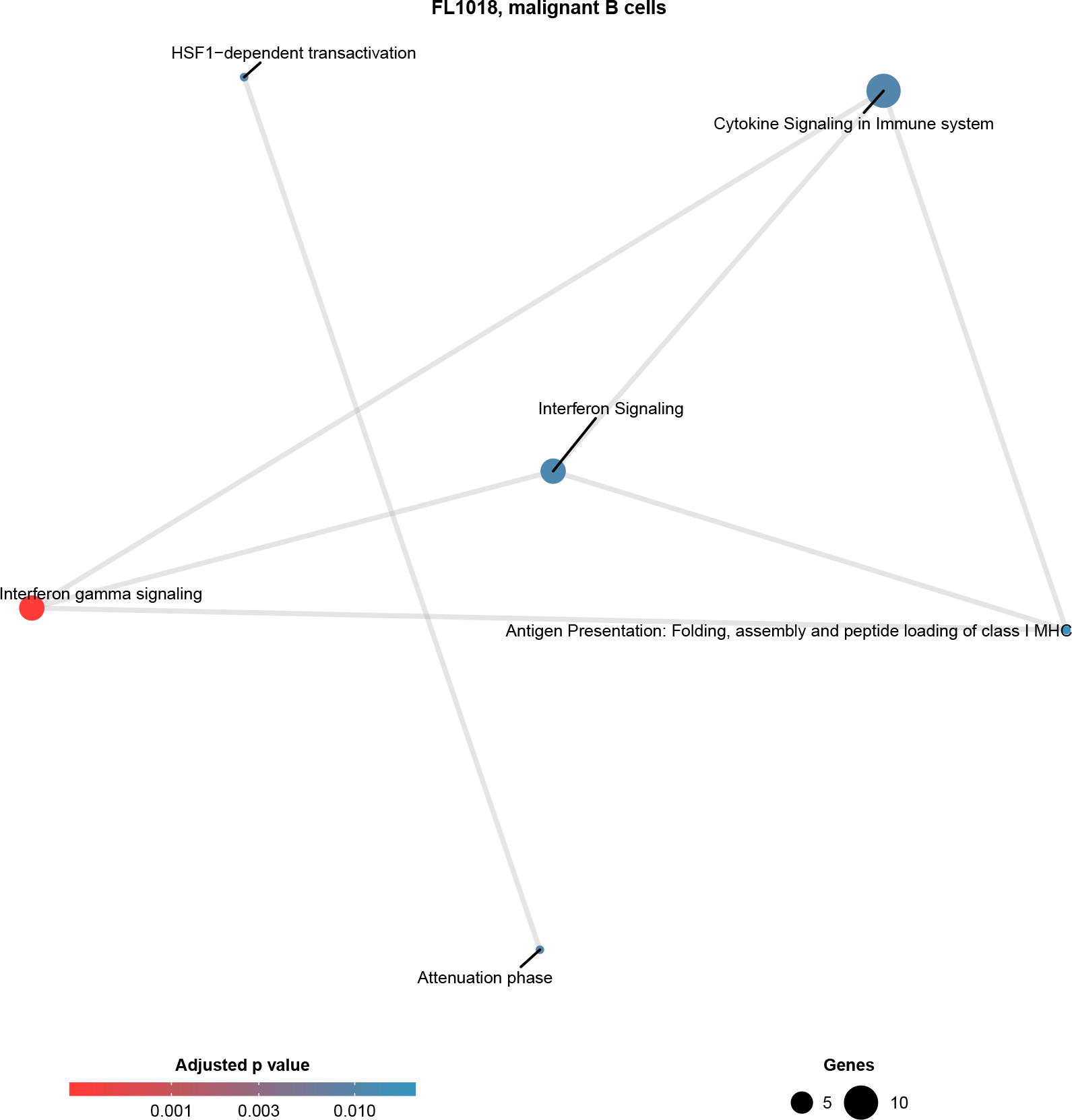
Significantly enriched Reactome pathways (BH-adjusted *P* -value ≤ 0.05) among the top 50 most downregulated genes (ranked by log fold change) in malignant B cells in FL1018. No pathways were significantly downregulated in FL2001.

**Supplemental Figure 17.**
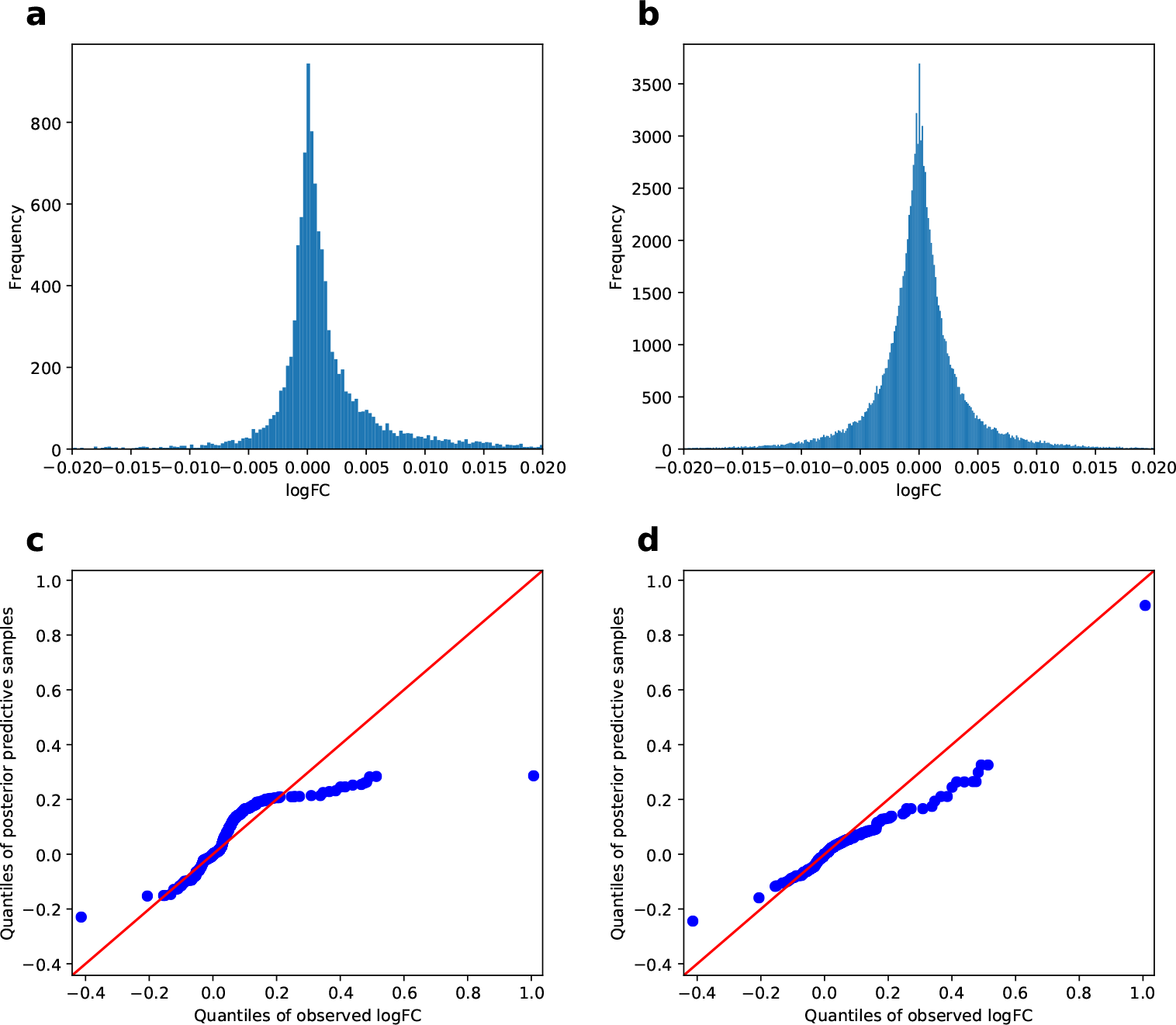
Fitting single cell RNA-seq simulation models to the Zheng PBMC 68k dataset, using cell type annotations provided in [38]. (**a**) Log fold change values computed from differential expression analysis between naïve CD8+ and naïve CD4+ T cells. (**b**) ‘Null’ log fold change values computed by randomly splitting naïve CD8+ T cells into equally sized halves 10 times. (**c**) Quantile-quantile (QQ) plot comparing observed log fold change values between naïve CD8+ and naïve CD4+ T cells and posterior predictive samples from the splatter model (**Methods**). (**d**) Quantile-quantile (QQ) plot comparing observed log fold change values between naïve CD8+ and naïve CD4+ T cells and posterior predictive samples from the modified model (**Methods**).

**Supplemental Figure 18.**
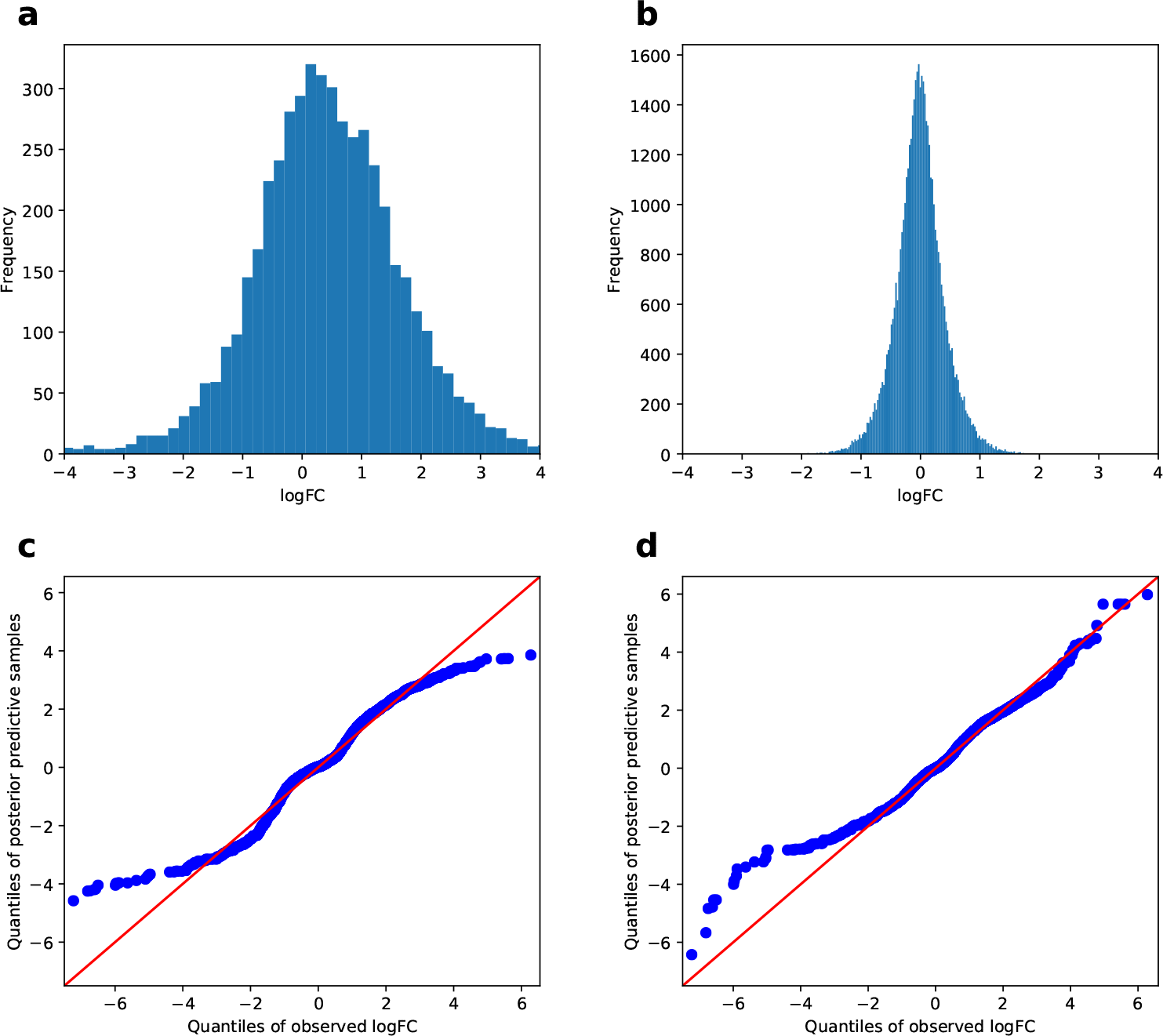
Fitting single cell RNA-seq simulation models to the Koh et al. [12] dataset of FACS-purified cell types. (**a**) Log fold change values computed from differential expression analysis between human embryonic stem cells (hESCs) and day 3 somite cells (ESMT). (**b**) ‘Null’ log fold change values computed by randomly splitting anterior primitive streak cells into equally sized halves 10 times. (**c**) Quantile-quantile (QQ) plot comparing observed log fold change values between hESC and ESMT cells and posterior predictive samples from the splatter model (**Methods**). (**d**) Quantile-quantile (QQ) plot comparing observed log fold change values between hESC and ESMT cells and posterior predictive samples from the modified model (**Methods**).

**Supplemental Figure 19.**
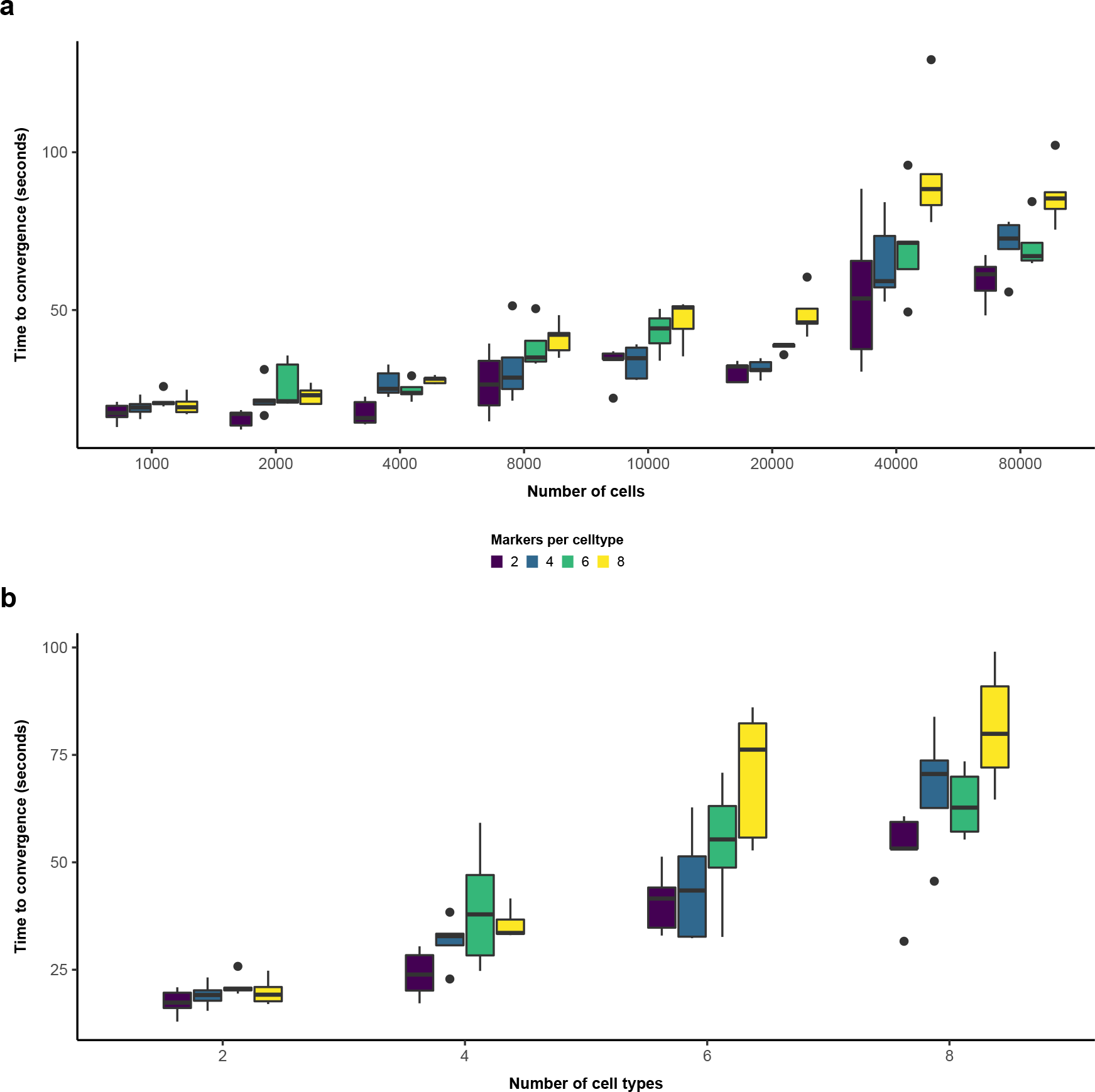
Benchmarking results for CellAssign across a range of simulated data set sizes (number of cells), number of cell types being inferred, and number of marker genes per cell type. (**a**) Runtime (to convergence, defined as a relative change in log-likelihood < 10^−3^ between successive iterations, as a function of data set size and the number of marker genes used per cell type, on simulated data (**Methods**). Two cell types were used. (**b**) Runtime (to convergence, defined as a relative change in log-likelihood < 10^−3^ between successive iterations, as a function of the number of cell types and the number of marker genes used per cell type, on simulated data. One thousand cells were used.

**Supplemental Table 1.** Performance measures on simulated data.

**Supplemental Table 2.** Marker gene matrices used in analysis.

**Supplemental Table 3.** Pathway enrichment results for follicular lymphoma and HGSC data.

